# traits.build: a data model, workflow and R package for building harmonised ecological trait databases

**DOI:** 10.1101/2024.02.11.579848

**Authors:** Elizabeth Wenk, Payal Bal, David Coleman, Rachael Gallagher, Sophie Yang, Daniel Falster

## Abstract

Trait databases have proliferated over the past decades, facilitating research on the ecology, evolution, and conservation of taxa across the Tree of Life. Typically, teams of independent researchers build these databases, and each must develop their own workflow and output structure. This divests research hours from downstream tasks such as trait-based analysis and interpretation and the resultant datasets are often difficult to integrate due to disparate database structures. Here we introduce the {traits.build} package, which offers a generalised workflow for building trait databases. {traits.build} contains bespoke functions for propagating metadata files, extensive tutorials, and sample configuration files, allowing researchers to efficiently build a new trait database using open source tools. In addition, the {traits.build} output structure is fully documented by a data model, ensuring the meaning of each variable and sematic relationship between variables is transparent and consistent. The data standard links to terms in previously published data standards, drawing strongly on DarwinCore and the Ecological Trait-data Standard, but also includes the ability to fully map location and context properties absent from these vocabularies., It is the first published database-building workflow that adheres to the Extensible Observation Ontology. Simultaneously developing a generalised workflow and publishing a data standard for the workflow provides {traits.build} users a straightforward pathway to build a new trait database that achieves the FAIR principles. The meaning of all variables in a {traits.build} database are already documented, allowing further integration with either other {traits.build} databases or indeed any other database with a documented data model. This follows the vision of the Open Traits Network to build trait databases whose data can be easily integrated for further analysis.

## Introduction

Trait data are central to an increasing array of ecological, evolutionary, and applied questions; informing our understanding of ecological strategies and evolutionary trajectories across the tree of life (Darling et al., 2017; Díaz et al., 2016; Meiri, 2018; Oliveira et al., 2016; Sauquet et al., 2017), assessments of biodiversity (Li et al., 2022; Vandewalle et al., 2010), management decisions (Harvey et al., 2022; Mayfield et al., 2010) and predictions of species responses to extreme events (Gallagher et al., 2022). For any given analysis, trait data are sometimes collected afresh, but, commonly, are also drawn from a multitude of local, regional, or global databases (Falster et al., 2021; Frimpong and Angermeier, 2009; Kattge et al., 2020; Madin et al., 2016; Myhrvold et al., 2015; Open Traits Network, 2023; Tobias et al., 2022). Indeed, as knowledge about the traits of an organism become known and available, it makes little sense to recollect this, just as one draws upon a previously established taxonomy or genome for an established species. Yet, despite the enormous growth and proliferation of trait databases during the last 20 years, the recording and organisation of trait data for the world’s biodiversity is still in its infancy. That is, despite major progress in some groups such as plants, we have little systematically recorded trait data on many branches of the tree of life, such as invertebrates and microorganisms (Gallagher et al., 2020). Compared to taxonomic and occurrence data, trait data lack established data models, a network of aggregators, workflows, and vocabularies. Moreover, trait data is arguably harder to organise; there is more information to collate and harmonise, traits are often linked and capturing these dependencies is hard, and the traits of interest usually differ among taxonomic groups. Researchers frequently find themselves attempting to parse complex data structures into a standardised format but lack the required tools and expertise. Workflows for building trait databases are therefore greatly needed.

A common challenge for aggregation of trait data, is that such data mostly arises in small heterogeneous fragments, which must later be reconciled and merged to create a database. Commonly, traits are collected for an individual scientific study or taxonomic review. As science proceeds paper by paper, trait data becomes available with similar granularity. Only once enough research papers have collected similar trait data across a variety of locations or species, can these be aggregated for further analyses, and then perhaps turned into or submitted to a database.

Moreover, these small data fragments have diverse formats, having been collected by diverse humans and in absence of established standardised formats. Big datasets in trait ecology therefore almost always come from amalgamation of many small, heterogeneous datasets, termed long-arm ecological datasets (Gerstner et al., 2017; Parr et al., 2016; Vanderbilt and Gries, 2021). Further, unlike data on species occurrences and taxonomy, there are not yet any widely implemented data models for data on traits. A data model – or data standard – is a description of the variables in a database and the relationship between variables, and, to the extent possible, these variables and relationships are matched to previously published terms (vocabularies) facilitating understanding and reuse of the data model. Without common workflows, every trait database that currently exists, has used a different data model and process to ingest and harmonise data, creating inefficiencies and challenges in data exchange.

Moreover, as most databases have been amalgamated to address specific research questions, their breadth, structure, and information content varies widely. Many trait databases report taxon-level means (Jones et al., 2009; Weigelt et al., 2020; Wilman et al., 2014), while others also record individual measurements (Kattge et al., 2020; Madin et al., 2016; Tobias et al., 2022; White et al., 2019); some output a single data table, while others are a collection of relational tables; and of course, they differ in the columns included and the naming conventions for these columns. Only a few databases record all contextual properties, despite a consensus that trait data can be more efficiently and reliably reused if the context surrounding each trait value is recorded (Farley et al., 2018; Kattge et al., 2011; Lenters et al., 2021; Madin et al., 2007; Vanderbilt and Gries, 2021). The challenge to accurately and completely document trait value metadata is amplified for databases that amalgamate many small datasets, each collected by a different researcher, documenting the metadata and context properties relevant to their research question.

The ecological trait database community increasingly recognises that all databases should be accompanied by an explicit data model, such that database users and the broader bioinformatics community are aware of how different pieces of information are documented and what assumptions are made (Kattge et al., 2011; Kissling et al., 2018; Lenters et al., 2021; Schneider et al., 2019). For the past decade, a universal message in papers discussing new ecological databases, new database standards, or the need for standards, emphasises how little standardisation exists in both how biologists circumscribe traits and how trait value metadata is documented (Garnier et al., 2017; Parr et al., 2016; Vanderbilt and Gries, 2021).

As identified by the informatics community, there are two different components to a database data model: 1) explicit descriptions of the variables, each database column header, whenever possible linking terms to previously published ontologies that include the same concept; and 2) a relational structure that describes how individual cells within the database are semantically linked. Ontologies are formal documentations of a data model, “a set of concepts and categories in a subject area or domain that shows their properties and the relations between them” (Oxford Living Dictionary) and may include just descriptions of each term or both term descriptions and the relationships between them. For instance, the ontologies for Darwin Core (DwC) (Wieczorek et al., 2012) and the Ecological Trait-data Standard (ETS) (Schneider et al., 2019) offer explicit descriptions of each term within each Class (table), with the ETS linking back to the identical or similar concept within DwC. Neither currently includes a relational data model, although DwC will shortly be releasing one (https://www.gbif.org/new-data-model). Meanwhile, the Extensible Observation Ontology (OBOE) (Madin et al., 2007) and the Sensor, Observation, Sample, and Actuator (SOSA) ontology (Janowicz et al., 2019) offer generic data models that describe the semantic relationships between pieces of information and can be incorporated into individual databases’ data models. The overheads for creating a suitable data model are large, hence few trait databases currently have a formalised and public data model.

Here we present the traits.build data model, workflow and R package, an open-source pipeline for compiling harmonised ecological trait databases from diverse sources. Traits.build began as the workflow for building the AusTraits plant trait database (Falster et al., 2021) and has since been expanded and generalised, establishing a database structure to support the creation of diverse ecological trait databases. The workflow is designed to bring together diverse datasets by aligning trait concepts and trait values, annotating trait values with all relevant metadata, then outputting the data in a standardised format documented by a published data model. It has, to date, been used for three compilations, covering disparate taxonomic groups and trait concepts. Simultaneously developing the workflow and using it to build varied databases has led to a workflow that is robust, straightforward to modify, and generalised to diverse use cases.

## Methods

The traits.build data model, workflow, output tables and R package were developed based on specified requirements (below), in an iterative process, where all components were gradually refined as we incorporated diverse datasets into three different data compilations.

### Database Requirements

The following requirements drove development of the materials.

#### Data model

1) Accommodates all metadata and contextual data associated with a trait measurement.
2) Follows best-practises outlined by the bioinformatics community, including:

a) Defines all variables in the output structure.
b) Documents the semantic relationships between database variables.
c) Uses previously published terms whenever possible.
d) Publishes the data model as a formal ontology, i.e. machine-readable format.

#### Output structure

3) Outputs are documented and fully adherent to the established data model.
4) Documents all metadata and contextual data associated with a trait measurement.
5) Given the large number of trait concepts and associated metadata, uses a relational structure and long data format, for efficient data storage.
6) Subsets of data can easily be extracted and/or manipulated, e.g. transformed into a wide format.
7) Different tables can easily be merged into a single unified table.

#### Workflow

8) Enables curators to combine diverse sources into a single harmonised, error-checked database with aligned trait names, units, categorical trait values, and taxon names.
9) Uses and leverages free and leading tools in data science, including R, GitHub.
10) Entails no cost to database creators/maintainers.
11) Is transparent and reproducible.
12) Enables the entire database to be rebuilt from source datasets at any time.
13) Enables diverse sources to enter a common pipeline, with minimal modification of original data sources.
14) Encourages data contributors to provide data “as is”, i.e. without restructuring into a specific format.
15) Enables database curators to revisit decisions made in the harmonisation of data sources.
16) Builds confidence for both dataset contributors and curators in the quality of outputs.

#### R package

17) Code is generalised into package format, so that it can easily be reused.
18) Package has extensive tests and automated testing to ensure it behaves as described.

### Open source tools

The workflow was built in R, one of the predominant coding languages used for data science and especially within the ecological sciences. As R is entirely open-source, there are no costs to using it, and all our code is entirely open-source, developing trust with the user community. We used GitHub for collaborative development of the tools, at the following open source repositories: https://github.com/traitecoevo/traits.build, https://github.com/traitecoevo/austraits.build.

Within R, we leveraged open source packages, including diverse packages from tidyverse (dplyr, tidyr, readr, stringr, lubridate, rlang, purrr) (Wickham et al., 2019), testthat (Wickham, 2011), and yaml (Garbett et al., 2023).

### Example compilations

We applied our methods to assemble the following three databases during the development process. Using the workflow and confronting it with diverse data sources, helped identify issues to generalise and improve the approach.

1. AusTraits: compiles plant trait data for Australia’s 30,000 native and naturalised plant taxa and includes some data for more than 500 traits (https://zenodo.org/records/10156222).
2. AusInverTraits: traits data for Australian invertebrates from 160 distinct sources, describing 53 invertebrate traits for 4654 taxa (unpublished, https://zenodo.org/records/10023667)
3. AusFizz: a plant physiology database housing response curve data and raw instrument output for Australian plant species. (unpublished, https://github.com/traitecoevo/AusFizz)

## Results

To date, more than 370 datasets spanning more than 1.8 million measurements have been merged into the AusTraits database, over 25000 measurements from 160 datasets into the AusInverTraits database, and more than 2000 response curves from 30 datasets into the AusFizz database, offering a broad span of datasets to fine-tune the traits.build data model, output structure, and pipeline.

Datasets are occasionally contributed to one of these databases that have a structure or metadata content that has not previously been encountered. However, iterations across > 5 years have led to a dataset input structure, database building pipeline, output structure, and data model that support all dataset formats submitted.

### Minimal ingredients for a database

The minimal database repository required to build a new database includes just four configuration files and a pair of data/metadata files for each data source.

### Configuration files

Four configuration files are required for the compilation of a {traits.build} database: 1) a trait dictionary; 2) a unit conversion file; 3) a taxon list; and 4) a database metadata file (Figure 1). Within the database repository, these folders are stored in the ‘config’ folder. Sample files are downloadable as part of the {traits.build} starting template, available at https://github.com/traitecoevo/traits.build-template/.

**Figure 1.**
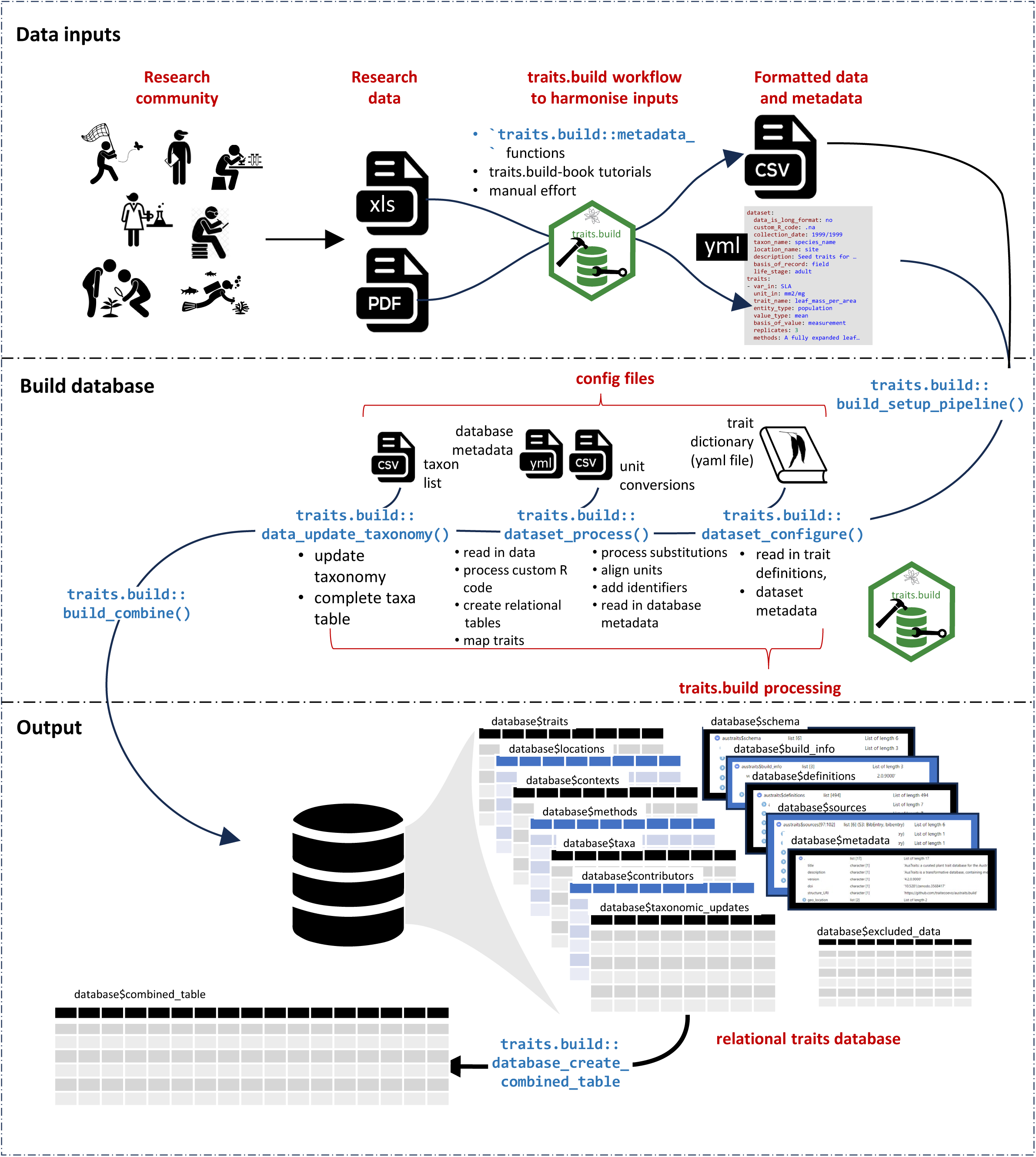
The traits.build workflow offers a pipeline to build harmonised trait database from disparate tabular data inputs provided by the research community. It includes functions to propagate a structured metadata file to accompany each dataset. It then includes an automated workflow that merges the datasets into a single database. A core component are three metadata files that specify the rules for harmonizing taxon names, trait concepts, and units. A four configuration file documents database metadata. The core database comprises thirteen relational tables, which can be combined into a single output table with a bespoke function.

#### Trait dictionary

A YAML-format file, traits.yml, is a list of defined trait concepts. Required fields for each trait include a label, a description, the trait type (categorical or numeric), accepted units (for numeric traits), an allowable range (for numeric traits), and allowable values (for categorical traits; must include a description of each trait value). Additional fields are allowed and encouraged; for instance, in the ‘traits.yml’ file for the AusTraits plant trait database, trait concepts each have a unique URI, referencing a trait concept published within the AusTraits Plant Dictionary (Wenk et al., 2023). The OntoPortal repositories BioPortal, EcoPortal and AgroPortal offer a selection of domain-specific published vocabularies to reuse (Jonquet et al., 2023).

#### Unit conversions

The unit conversions.csv file will generally be interchangeable across databases and a copy of the AusTraits unit conversion file is included in the {traits.build} starting template. It has three columns: the input units (units corresponding to the trait data values in the input data table), the accepted units for a specific trait concept, as indicated in the trait dictionary, and the numeric conversion required to convert from the input to the output units. For instance, a row might read: ‘m,mm,x*1000’, indicating 1 metre is equivalent to 1000 millimetres. It is best practise to consistently use unit abbreviations specified in a published vocabulary, such as those defined by the Unified Code for Units of Measure (UCUM; https://ucum.org/ucum)(Schadow and McDonald, 2017).

#### Taxon list

The taxon_list.csv file documents information about the taxa represented in the database, including taxon names, taxon ranks, taxonomic status, higher order taxon groupings, and taxon identifiers. The information must be independently compiled based on the available taxonomic resources for the taxon groups covered in a specific trait database. To accommodate diverse databases and the varied taxonomic resources available across the Tree of Life, the {traits.build} pipeline requires only three columns, ‘taxon_name’, ‘aligned_name’, and ‘taxon_rank’, but it is best practice to also include taxon identifiers and higher-order groupings (e.g. genus, family, etc.)(Figure 1). The ‘aligned_name’ is intended to be a scientific name where typos have been corrected, but which might not yet have been updated to the name of a currently accepted/valid taxon concept. The ‘taxon_name’ is then the best-matched name for a taxon; it will most commonly be the name of an accepted/valid species or infraspecific but may include informal names if an accepted/valid name is not available.

#### metadata.yml

This is a YAML-format file that documents database metadata, including the database name, creators, licence, and version. Fields within the ‘metadata.yml’ file include all those considered essential by Datacite 4.4 (2021) to make a data resource FAIR (Wilkinson et al., 2016). A mostly empty version of the file is available as part of the {traits.build} starting template.

Information that is identical for all {traits.build} databases is already filled in, such as specifying that the database is compiled using the {traits.build} R package.

### Dataset files

Inputs are organised by source, with a folder for each data source assigned that dataset’s ‘dataset_id’, standardly in the format ‘Contributor Surname_Year’. Within this folder are two required files, ‘data.csv’ and ‘metadata.yml’. The ‘data.csv’ file is a spreadsheet containing all trait data, taxon names, location names (if relevant), and any context properties (if relevant). The ‘metadata.yml’ file assigns the columns from the ‘data.csv’ file to their specific variables and maps additional dataset metadata in a structured format (Supplementary Table 1). Within the database repository, these folders are stored in the ‘data’ folder.

### Training resources

Training resources have been developed to encourage the use of the {traits.build} R package. These are hosted on two accompanying GitHub repositories,

- https://github.com/traitecoevo/traits.build-template: Includes three resources 1) sample configuration files that can be used as a starting point for a new database; 2) two example datasets (‘example_dataset_1’ and ‘example_dataset_2’) that include both datasets and propagated metadata files to trial the building of a simple {traits.build} database; and 3) multiple tutorial datasets to practice propagating metadata files.
- https://github.com/traitecoevo/traits.build-book: A complementary resource, including tutorials to accompany each of the tutorial datasets, with instructions on how a combination of functions and manual editing can be used to map diverse dataset formats into dataset ‘metadata.yml’ files. This repository also offers detailed vignettes about the {traits.build} database structure, troubleshooting, and how to explore and wrangle data in a {traits.build} database.

### Data model and ontology

Following a review of available relational ontologies, the traits.build database structure was developed to be compliant with the OBOE data model (Madin et al., 2007). OBOE offers an observation-centric modular ontology developed explicitly for documenting the complexity of ecological data. In OBOE, each ‘observation’ encompasses ‘measurements’ of multiple traits made at a single point in time (Figure 2). Each observation is of a single ‘entity’, where the ‘entity’ for trait measurements is most commonly an individual organism, population, or species. The ‘measurements’ are of a ‘characteristic’ (a trait) and each reference a specifically described ‘standard’, a trait concept defined in an accompanying resource, the traits dictionary in this case.

**Figure 2.**
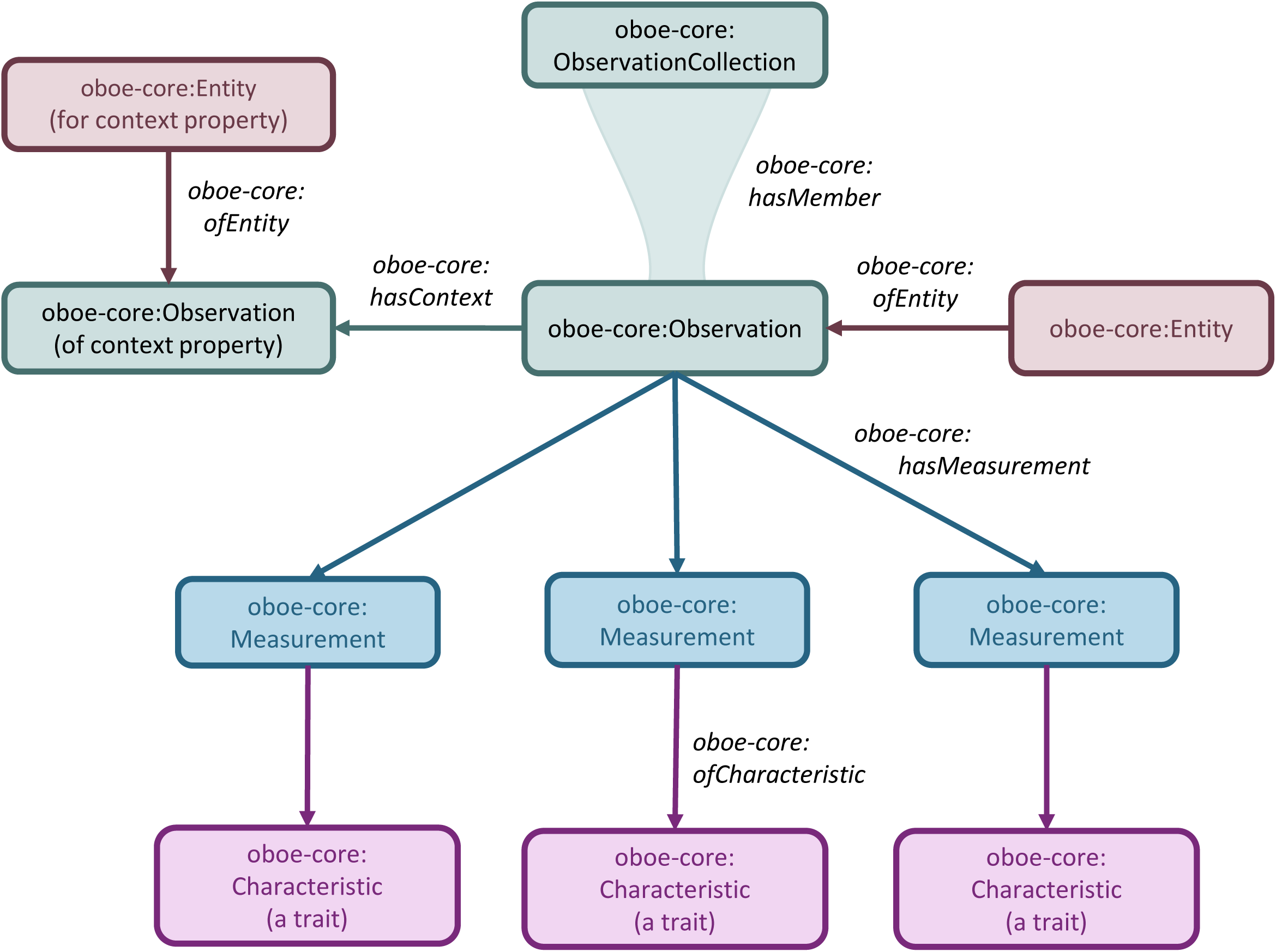
The core Extensible Observation Ontology (OBOE) data model. The core classes in OBOE are : 1) an **entity** (the object of interest, such as an individual plant); 2) an **observation** of the entity, including all measurements made on a single entity at a single point in time; 3) a **measurement** which is a single trait measurement that is part of an observation; and 4) a **measured characteristic**, which is the property being measured (a specific trait). Groups of observations can be grouped into **observation collections**, such as observations of multiple individuals made at a specific location. In addition, the **has context** property asserts that one observation can be a context for a second observation, suggesting that the observation of a context property helps with the interpretation of the measurements that comprise the core observation.

The OBOE data model also links ‘observations’ of context properties to ‘observations’ made on biological entities. Just as an individual organism is an ‘entity’, so is a location an ‘entity’, and ‘measurements’ can be made of location properties. By linking these to ‘observations’ of the biological organism, contextual information can be mapped into the data model. Finally, ‘observations’ can be grouped together into ‘observation collections’ that share, for instance, a common context property, location, or taxon.

Using the OBOE ontology allowed the development of a data model into which ecological trait measurements and all accompanying context properties could be mapped with semantic clarity (Figure 3). For all measurements of organismal traits, location properties, or context properties, there is always a clearly defined entity (individual organism, population, taxon, location, treatment context, etc.) which has been observed (Table 1, Figure 3). One observation can offer context for another observation (or observation collection), such as an observation of a location offers context for an organismal trait observation at that location. Observations of context properties are assigned a specific category: treatment contexts, plot contexts, entity contexts, and temporal contexts (Tables 1, S2, Figure 3).

**Figure 3.**
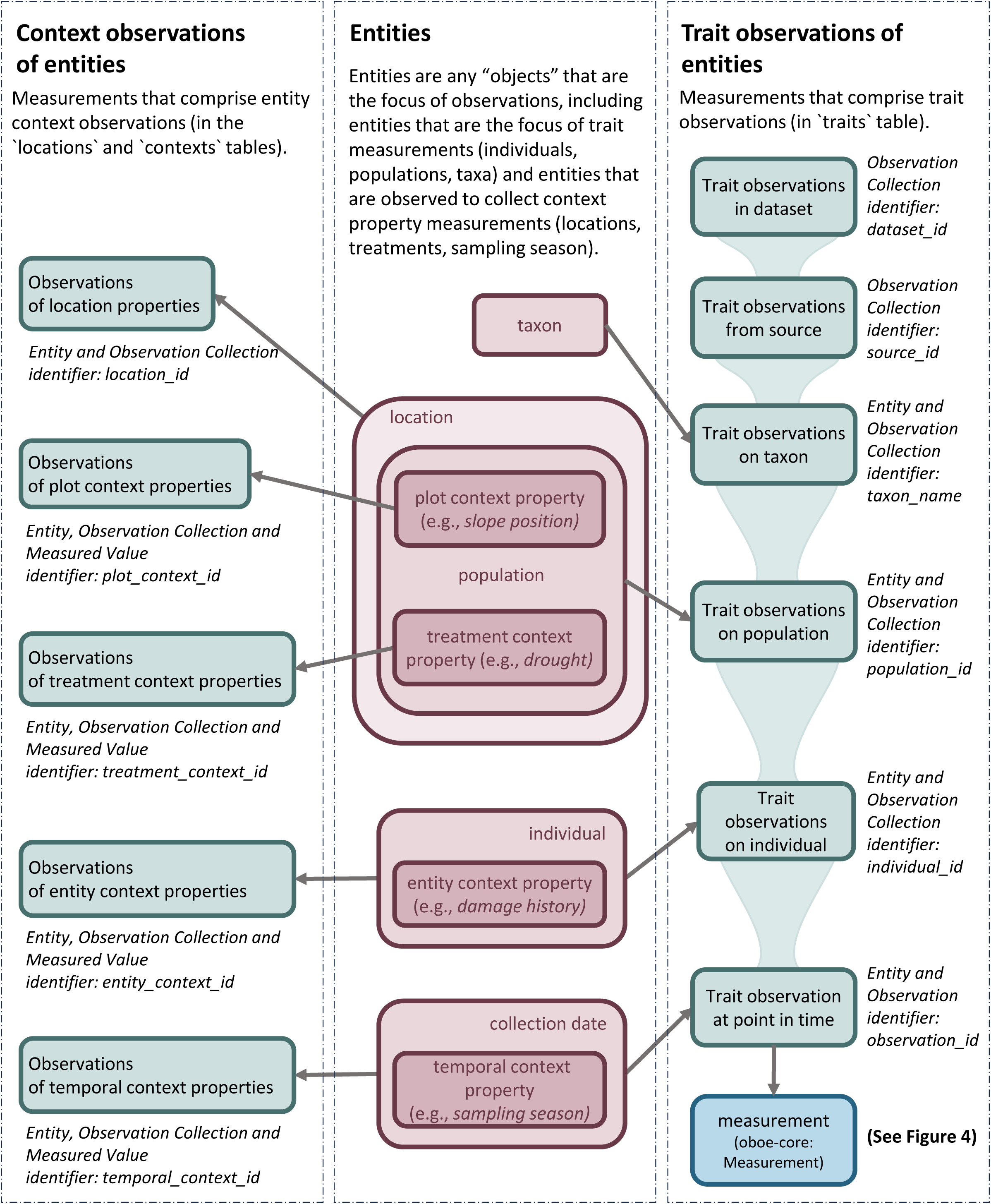
Within the traits.build output tables, a collection of identifiers uniquely identify specific entities, observations, and observation collections. Observations of entities may yield both trait measurements and context property measurements. For instance, an individual may be observed both to measure trait values (i.e. leaf area and leaf thickness) and context properties (i.e. damage history or canopy position). Within traits.build, a population is defined as a collection of individuals from the same taxon growing in the same location <u>and</u> experiencing the same contextual situation (i.e. same plot context and treatment context values.)

**Table 1.** Specifically identified observations and observation collections within a traits.build database and the links between observations/observation collections and entities, measurements and identifiers that are specified in the {traits.build} data model.

| Observation /<br>observation collection | Description | associated<br>Entity | associated Measurement(s) | identifier | identifier links to |
| --- | --- | --- | --- | --- | --- |
| Observation | An Observation is a collection of Measurements made at a single point in time. In a {traits.build} database it is the subset of rows within the traits data that are made on the same entity and share the same observation_id. | Entity (any) | trait measurements that comprise an observation | observation_id | Observation |
| observations from source | Observation collection encompassing all observations of organismal traits, location properties, and contexts for a single published or unpublished source. | source | measurements linked to a single source | source_id | all measurements in observation collection |
| observations of taxon | Observation collection encompassing all observations on a single taxon within a single dataset. | taxon | measurements linked to a single taxon | taxon_name | all measurements in observation collection |
| observations at location | Observation collection of location properties and organismal traits at a single location. | location | measurements m | location_id | all measurements that comprise observation |
| observations of population | Observation collection encompassing all observations on a single population. | population | Observation Collection | population_id | all measurements in observation collection |
| observations of individual | Observation collection encompassing all observations on a single individual. | individual | Observation Collection | individual_id | all measurements in observation collection |
| observation of treatment context | Observation where the entity is an experimental manipulation that might affect the trait values measured on an individual, population or species-level entity. | treatment context | measurement of treatment context property | treatment_context_id | individual measurement |
| observation of plot context | Observation where the entity is a plot (subset of a location) that might affect the trait values measured on an individual, population or species-level entity. | plot context | measurement of plot context property | plot_context_id | individual measurement |
| observation of entity context | Observation that records information about an organismal entity (individual, population or taxon) that does not comprise a trait-centered observation but might affect the trait values measured on the entity. | individual, population or taxon | measurement of entity context property | entity_context_id | individual measurement |
| observation of temporal context | Observation where entity is a category of context indicating repeat observations have been made on a plot, which might affect the trait values measured on individual or population-level entities. | temporal context | measurement of temporal context property | temporal_context_id | individual measurement |
| observation of method context | Observation and accompanying measurement that records specific information about a measurement method that is modified between measurements. | method context | measurement of method context property | method_context_id | Measurement |
| subset of an observation | A unique measurement within a measurement sequence that documents a response curve. | individual | trait measurement comprised of repeat measurements | repeat_measurements_id | individual point in a measurement |

The OBOE framework also relates each ‘measurement’ to a specific ‘characteristic’ and ‘standard’ (Madin et al., 2007) (Figure 4). A ‘characteristic’ is the trait that has been measured, while the ‘standard’ is a specific trait concept described within a standalone resource. In the {traits.build} data model, the ‘characteristic’ is indicated by the ‘trait_name’ within the traits table, while the ‘standard’ is the trait concept described within the traits dictionary file (and preferably linked to a published trait concept) (Figure 4). Finally, each measurement has a ‘measured value’ (the trait value). Figure 4 also depicts additional measurement metadata documented by the {traits.build} data model. For instance, metadata associated with individual measurements includes the method and method context, the latter indicating any modifications to the method stratified across individual measurements that are likely to affect the trait value. Additional measurement metadata includes two dataset-level fields, the dataset description and sampling strategy (Figure 4). There is also metadata associated with individual trait values (Figure 4).

**Figure 4.**
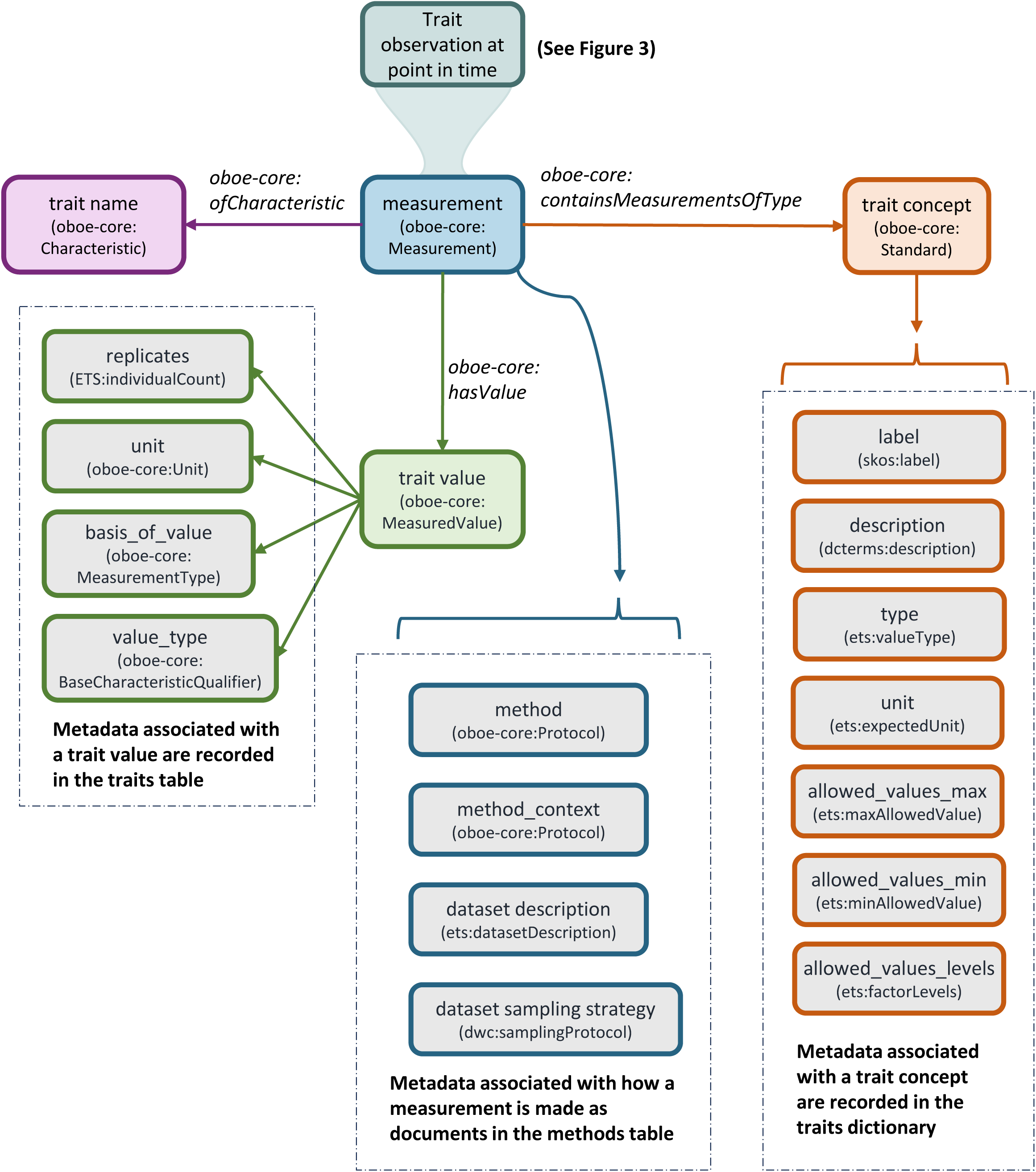
The traits.build workflow ensures that all metadata associated with a trait concept, a trait measurement, and a measured value is documented. The trait concept metadata is documented in the traits dictionary, a yml-formatted file that includes a trait description, trait type (numeric versus categorical), allowed categorical trait values, minimum and maximum allowed values (for numeric traits), expected unit (for numeric traits), and a trait label (its “name”). Metadata associated with a measured trait value includes the units, the number of replicates, the value type (raw value, mean, minimum, maximum, mode, bin), and the basis of value (measurement, expert score, model-derived). This information is “context” for the trait concept, measurement,

These terms and other dataset and measurement metadata documented in the database’s relational tables were explicitly defined and matched to terms in previously published domain-specific vocabularies whenever possible (Table 2, S3-S7). In particular, DwC (Wieczorek et al., 2012) was selected as the authoritative ontology for classes documenting taxonomic information. For trait-related fields, the {traits.build} data model aligns predominantly with the terms defined by ETS (Schneider et al., 2019) (Figure 4), with additional best-practice fields to include defined within the AusTraits Plant Dictionary (Wenk et al., 2023). Additional published vocabularies used in the traits.build data model include DataCite 4.4 (https://schema.datacite.org/meta/kernel-4.4/) (2021) for fields documenting metadata relating to dataset contributors, and BibTex (https://zeitkunst.org/bibtex/0.2/) for fields relating to dataset references. In addition, classes from the Semanticscience Information Ontology (SIO) (Dumontier et al., 2014), Simple Knowledge Organization System (SKOS; https://www.w3.org/TR/skos-primer/) (Miles and Bechhofer, 2009), Resource Description Framework (RDF; https://www.w3.org/TR/rdf11-primer) (Brickley and Guha, 2014), and Dublin Core Metadata Initiative (dcterms) (“DCMI Metadata Terms,” 2010) vocabularies were used.

**Table 2.**
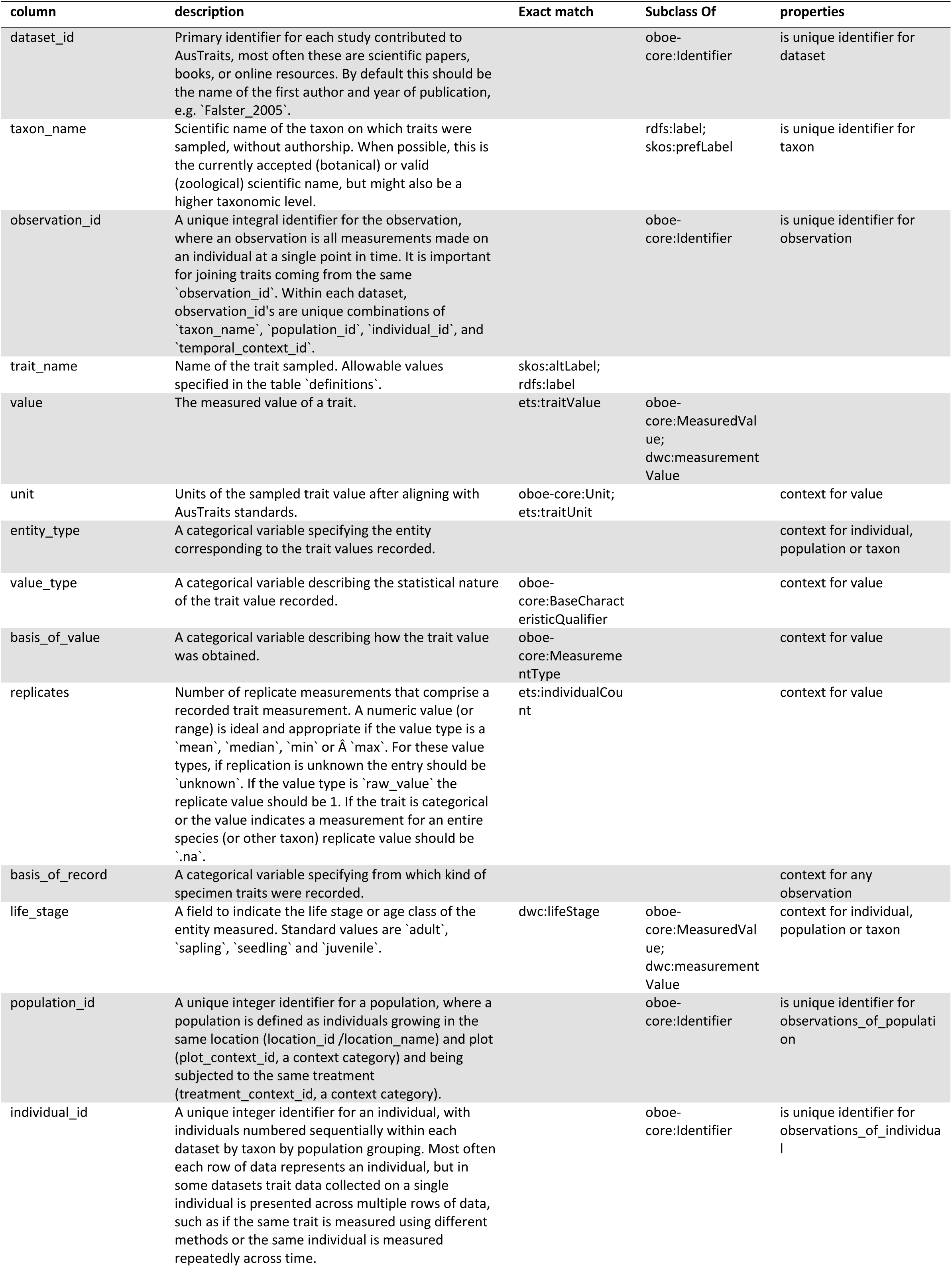
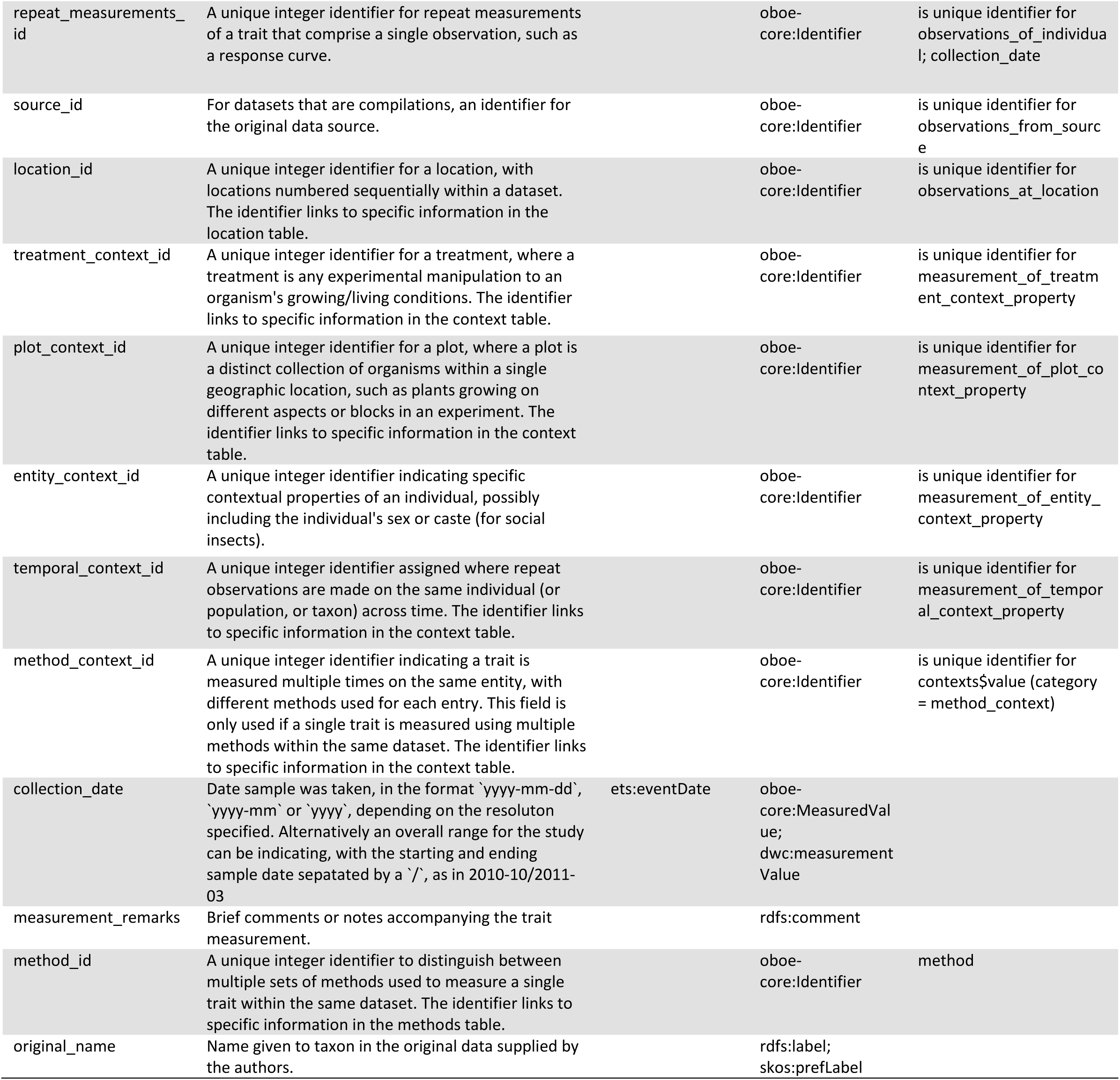
Columns within the traits table and their core mappings in the {traits.build} data model. See additional tables in the Supplementary Information for the other relational tables in a {traits.build} database.

The formal ontology or knowledge graph for {traits.build} therefore documents 1) the database structure’s adherence to OBOE; 2) descriptions for each of the terms (columns) output in a {traits.build} database; 3) links between {traits.build} terms and identical terms in published vocabularies; and 3) the relationships between individual columns, extending beyond those that can be documented using OBOE’s properties (Figures 3, 4; Tables 1, 2, S3-S7). The categories of information documented about each term are called annotation properties in formal vocabularies. Some annotation properties, such as a term description, term label, and term datatype were propagated for all terms (Table S8). Matches to identical, similar, or related terms within other vocabularies (skos:exactMatch; skos:closeMatch and skos:relatedMatch) and indications of superclasses (skos:broader) were made whenever possible. The remaining annotation properties are terms from either OBOE or the Semanticscience Integrated Ontology (SIO) that effectively capture the relationships between individual {traits.build} terms (Table S8). This information was documented in a tabular format, wherein there were rows for all {traits.build} terms and columns for all annotation properties; see https://github.com/traitecoevo/traits.build/tree/develop/ontology/traits.build_ontology.csv.

R scripts were used to wrangle the information from the table into a triples format, the core unit of the Resource Description Framework (RDF) data model (World Wide Web Consortium, 1999). With the triples format, all information content is collapsed into a single long-format document with three columns, the subject, the predicate, and the object. The subject is always the URI for a concept. For traits.build these are resolvable identifiers within the ‘https://w3id.org/traits.build’ namespace generated for each traits.build-defined term. The predicate indicates a property of the object that can be described; these are the annotation properties in Table S8 and each is represented by its URI. The object is the value for the specific predicate for the specific object and can be either a URI (for annotation properties that link to a previously published term) or a text string (for a term description). The R package {rdflib} (Boettiger, 2023) was used to output four separate RDF serialisations, Turtle (traits.build.ttl), N-Triple (traits.build.nt), N-Quad (traits.build.nq) and JSON Linked Data (traits.build.json) formats. Both these machine-readable and a human-readable html output of the data model are available at https://traitecoevo.github.io/traits.build/ontology/index.html. Individual terms are all documented on this page, or can be accessed via their individual URI’s, such as, w3id.org/traits.build/trait_name.

### Database structure

A {traits.build} database can be output in two distinct formats, the collection of relational tables generated by the R workflow (the primary output) or as a single combined table (Figure 1). The R pipeline creates a core traits table, supported by a series of tables documenting metadata (locations, contexts, methods, taxa, taxonomic updates, and contributors) (Tables 2, S3-S7). All tables are in long format, such that each trait measurement, location property measurement, context property measurement, etc. is recorded in a single row. This database structure is space-efficient, as each unit of information is stored only once. For instance, a location property measurement is recorded once in the location table, although many trait measurements will be linked to the bespoke location and its property. Similarly, the methods used by a specific study to measure a trait appear just once in the methods table. In the combined table, traits remain in long format, but the ancillary tables are joined in so that all relevant information for a single trait measurement appears in a single row.

The {traits.build} relational tables reflect that database’s adherence to the OBOE ontology (Figure 5). A single row within the traits, locations, and contexts tables is nearly always a single measurement; the exception is for trait concepts that are response curves of many values, the individual steps (sub-measurements) within the response curve are each recorded in a separate row. In the traits table, an observation consists of one or more traits measured on a single organismal entity at a single point in time. Similarly, within the location table, the multiple location properties measured at a single location comprise an observation. Observation collections can exist either within a single table or across the relational tables (Table 1, Figure 3). Identifiers are linked to unique observations, certain observation collections, or, for context properties, individual measurements (Table 1, Figures 3, 5). For instance, within a dataset a unique ‘observation_id’ is assigned to the measurements made as part of a single observation of an organismal entity. In addition to documenting the database’s relational structure, the identifiers allow the individual tables to be joined into a wider format.

**Figure 5.**
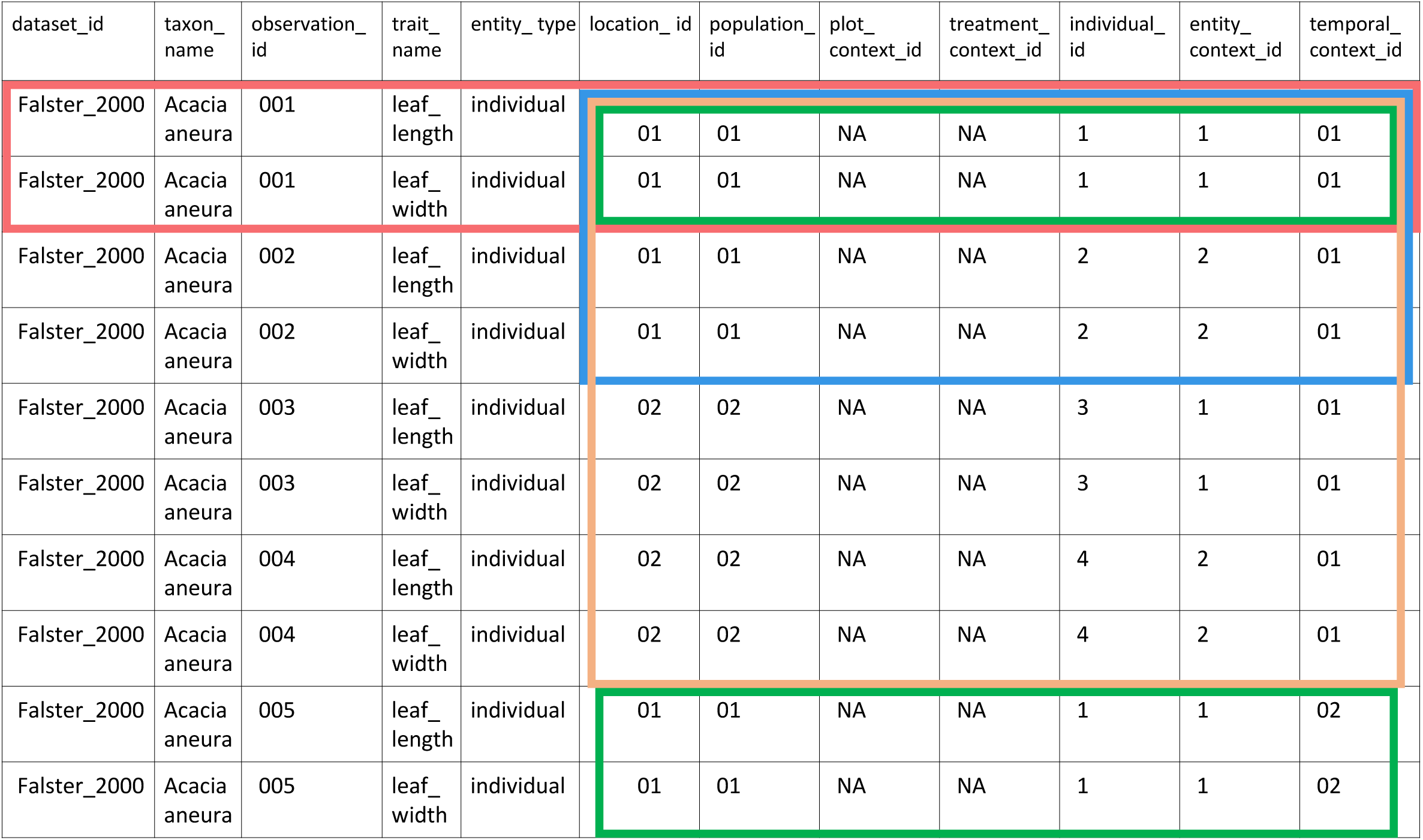
The traits table shows how the ‘{traits.build}’ output structure adheres to the OBOE data model. In this example, the Entity is an individual plant, growing in a specific location. Identifiers are used to link trait Observations to Observations of locations or contexts: 1) Measurements made on a single Entity at a single point in time comprise a single Observation and share an ‘observation_id’ and all other identifiers (----); 2) Measurements made on two individuals in the same location, at the same time, are automatically assigned separate ‘individual_id’ values. In this example, there is an additional entity context property observation linked to the individuals, such as, for instance, whether the individuals are ‘damaged’ or ‘undamaged’ by a disturbance (----); 3) Measurements are made on two individuals at each of two locations. Each individual is assigned a separate ‘individual_id’ values. In this example, individual 1 (at location 1) and individual 3 (at location 2) share a common entity context property value (e.g. ‘damaged’), so their ‘entity_context_id’ is identical (----); and 4) If the same individual is observed a second time, in conjunction with an observation of a temporal context property (i.e. season), the two sets of measurements will have different observation_id’s and different temporal_context_id’s, representing, respectively two observations of the individual (an entity) and two observations of season (an entity) (----).

Indeed, the combined table is a derived output, where the information in the seven relational tables is joined to the traits table. Measurements of location and context properties are collapsed into five compacted columns, ‘location_properties’, ‘treatment_context_properties’, ‘plot_context_properties’, ‘entity_context_properties’, ‘temporal_context_properties’, and ‘method_context_properties’. The trait measurements are still presented in long-format, but the table now includes 66 columns to include all dataset metadata in a single table. This format necessarily includes some duplication of data in these additional columns, greatly increasing the size of the resultant table.

### Pipeline to add datasets

Our primary goal with the workflow is to get different sources into a common pipeline, where each source is reshaped into a common format, described by the data model. The R-scripted workflow was designed to do this with original sources retained as close as possible to their original format and the decisions made in mapping the data into the database fully exposed and mutable. As identified by Lenters et al. (2021), to avoid translation errors automated processes are preferable for merging a researcher’s contributed spreadsheet into the database.

As indicated above, a folder is created for each data source within the database. Two files must exist within the folder, ‘data.csv’ and ‘metadata.yml’. These must be completed before the R-scripted pipeline can build (or rebuild) the database.

Building dataset data.csv files

All trait data for each dataset must be in a single spreadsheet, the dataset’s ‘data.csv’ file. If data for a study are submitted across multiple spreadsheets, these must be compiled into a single table. The {traits.build} workflow does not prescribe a specific process, but it is considered best practice to create a ‘raw\’ subfolder within the dataset’s folder where raw data files and code used to merge them are archived. Beyond combining multiple spreadsheets into a single ‘data.csv’ file, the ‘data.csv’ file should be minimally edited, with column headers, taxon names, and location names remaining identical to those in the data file contributed by the dataset custodian. Further details are available at https://traitecoevo.github.io/traits.build-book/adding_data_long.html#standardised-input-files-required.

### Building dataset metadata.yml files

The ‘metadata.yml’ file documents both what information is tabulated in each column in the ‘data.csv’ file and additional dataset metadata. The YAML-format metadata file that accompanies each ‘data.csv’ file is created using a combination of {traits.build} functions and manual input. YAML is a popular data structure for configuration files as it supports flexible nested information content and is easily human readable. A specific section of the ‘metadata.yml’ file allows the insertion of customised R code, to record manipulations that are performed to the ‘data.csv’ file as it is read into the R workflow. See https://traitecoevo.github.io/traits.build-book/tutorial_datasets.html for tutorials and examples.

#### Functions

Fifteen functions were developed for propagating the metadata.yml files (Table 3). These include functions to first render the skeletal ‘metadata.yml’ file (‘metadata_create_template()’), then to propagate information into the core sections of the ‘metadata.yml’ file (source, locations, contexts, traits, substitutions, taxonomic updates, and excluded data).

**Table 3.** {traits.build} functions to create and propagate information in a dataset’s metadata.yml file.

| function | required and optional user inputs | description |
| --- | --- | --- |
| metadata_create_template | dataset_id | Function to generate skeletal `metadata.yml` file with place-holders for major metadata sections. |
| metadata_add_source_bibtex | dataset_id, file, type = "primary" | Function to add source details to a dataset's `metadata.yml` file from a bibtex citation file. |
| metadata_add_source_doi | dataset_id, doi, type = "primary" | Function to add source details to a dataset's `metadata.yml` file from a doi. |
| metadata_add_locations | dataset_id, location_data | Function to add location properties to a dataset's `metadata.yml` file from a dataframe. The function asks users which columns in a wide-format metadata frame include location properties and records this appropriately in the metadata file. Manual editing may be required to change location property labels. |
| metadata_add_contexts | dataset_id, overwrite = FALSE | Function to add context properties to a dataset's `metadata.yml` file. The function asks users which columns in the dataset's `data.csv` file include context properties, the category of each context property, and whether value replacements and descriptions are required for each context property. The information is recorded in the metadata file, but manual editing is required to add context property labels, value replacements, and value descriptions. |
| metadata_add_traits | dataset_id | Function to add traits to a dataset's `metadata.yml` file. The function asks users which columns in the dataset's `data.csv` file include trait values and adds a template for those traits in the metadata. This template must then be finished manually. |
| metadata_add_substitutions | dataset_id, trait_name, find, replace | Function to align a single categorical trait value used in a dataset's `data.csv` file to the allowed categorical values for the specified trait. Allowed categorical trait values are those included for the trait in the `config/traits.yml` file. |
| metadata_add_substitutions_list | dataset_id, substitutions | Function to align a dataframe of categorical trait values used in a dataset's `data.csv` file to the allowed categorical values for the specified trait. Allowed categorical trait values are those included for the trait in the `config/traits.yml` file. |
| metadata_add_taxonomic_change | dataset_id, find, replace, reason, taxonomic_resolution, overwrite = TRUE | Function to add a single taxonomic change to a dataset's `metadata.yml` file. |
| metadata_add_taxonomic_changes_list | dataset_id, taxonomic_updates | Function to add a dataframe of taxonomic changes to a dataset's `metadata.yml` file. |
| metadata_update_taxonomic_change | dataset_id, find, replace, reason, taxonomic_resolution | Function to update a taxonomic change in a dataset's `metadata.yml` file. |
| metadata_remove_taxonomic_change | dataset_id, find | Function to remove a taxonomic change in a dataset's `metadata.yml` file. |
| metadata_find_taxonomic_change | find, replace = NULL, studies = NULL | Function to find all datasets whose `metadata.yml` files include a taxonomic change for a specific taxon name. The function can just search for matches to `original_names` (find) or also to `aligned_names` (replace). |
| metadata_exclude_observations | dataset_id, variable, find, reason | Function to exclude specific observations from a dataset, selecting observations by any variable and value (find). |
| metadata_check_custom_R_code | dataset_id | Function to check the output of running the `custom_R_code` specified in the `metadata.yml` file for a specified `dataset_id`. The function reads in the file `data.csv` for the specified `dataset_id` and applies manipulations included under `custom_R_code` in the file `metadata.yml`. |

#### Manual completion

After functions are used to add traits, locations, contexts, substitutions, and taxonomic updates to the ‘metadata.yml’ file, manual steps are required to complete all fields within the YAML file and edit the pre-populated fields where necessary. For instance, dataset contributors, some dataset and measurement metadata variables (i.e., life stage, sampling strategy), and the details for each trait must be filled in manually. The dataset curator must match each column that contains trait data to a specific trait concept from the accompanying trait dictionary and add units, value types, entity types, replicate counts, and methods for each trait.

#### Custom R code

The ‘custom_R_code’ section of the ‘metadata.yml’ file offers an opportunity to make additional manipulations to the ‘data.csv’ as it is read into the {traits.build} processing pipeline. Common uses for ‘custom_R_code’ are to replace placeholder symbols with NA’s (i.e. other characters, zeros, etc.), to replace duplicate values with NA’s (i.e. a species-level trait value that is reported as part of many individual-level observations), and to add a column with a location name (when the data.csv file lacks a column documenting the location name). The function ‘metadata_check_custom_R_code()’ offers a look at the ‘data.csv’ file to ensure the ‘custom_R_code’ manipulation has changed the ‘data.csv’ file as anticipated.

### Building the database

Guided by the information in the four configuration files (trait dictionary; units conversion file; taxon list; and database metadata file), the R-scripted workflow combines the ‘data.csv’ and ‘metadata.yml’ files for the individual datasets into a unified, harmonised database.

Before starting the build workflow, ‘build_setup_pipeline()’ is run to set up the build process, selecting from different build methods, choosing which datasets to build, and naming the output database (Figure 1). The database is then built (and rebuilt) with a single function line of code, ‘source(“build.R”)’ for the default build method. There are three distinct steps to the build process, processed by a trio of functions, ‘dataset_configure()’, ‘dataset_process()’, and ‘dataset_update_taxonomy()’ (Figure 1). These functions cyclically build each dataset, only combining them into a single database at the end of the workflow.

#### Configuring files (‘dataset_configure’)

During the first step, the metadata file is read in, and the suite of trait concepts required to process this dataset are subsetted from the ‘traits.yml’ file (Figure 1).

#### Processing each source (‘dataset_process’)

The core dataset processing begins by reading in the dataset’s ‘data.csv’ file and applying any ‘custom_R_code’ (Figure 1). It is followed by steps that use the rules documented in the dataset’s ‘metadata.yml’ file to process the ‘data.csv’ file, identifying which columns contain data pertaining to traits, taxon names, location names and contexts. This information is used to propagate a series of relational data frames, one for the core traits data, and additional dataframes that document metadata/measurements that pertain to locations, contexts, methods, data contributors, and taxonomic updates. An important component of ‘dataset_process’ is adding identifiers, e.g. location_id, observation_id, etc (Figure 1), that ensure the metadata compiled into ancillary tables remains linked to the corresponding trait observations. At this processing stage, an interim taxa dataframe simply documents all taxon names within the dataset.

A core flexibility coded into this step of the {traits.build} workflow is to allow values for some variables to be read in as either a fixed value or from a column in the ‘data.csv’ file. The fields ‘entity_type’, ‘value_type’, ‘basis_of_value’, ‘replicates’, ‘collection_date’, ‘unit_in’, ‘basis_of_record’, ‘life_stage’, ‘measurement_remarks’, ‘source_id’, and ‘methods’ can be input as a fixed value at the dataset (or trait or location) level, or have distinct values across rows in the ‘data.csv’ file. The R-workflow checks whether the value coded into the dataset’s ‘metadata.yml’ file is a column within the ‘data.csv’ file; if yes, the values in the column are read in; if no, the fixed value is used for all observations.

For the traits table there are a series of additional manipulations to ensure trait values are harmonised across datasets. Units are aligned to those specified for each trait concept in the ‘traits.yml’ file and numeric trait values are adjusted accordingly using the conversion factor specified in the ‘unit_conversions.csv’ file. Categorical trait values specified in the ‘substitutions’ section of the metadata file are applied, ensuring trait values are aligned to those in the traits dictionary. Any changes to the submitted taxon names that are specified in the ‘taxonomic_updates’ section of the ‘metadata.yml’ file are made. Trait measurements are flagged for exclusion if they are either explicitly listed in the ‘exclude_observations’ section of the ‘metadata.yml’ file or if trait values are unacceptable based on a series of checks against the trait dictionary. Possible reasons for excluding data include: “Observation excluded in metadata”, “Trait name not in trait dictionary”, “Unsupported trait value” (for categorical traits), “Value does not convert to numeric” (for numeric traits), “Value out of allowable range” (for numeric traits), “Missing unit conversion” (for numeric traits), “Missing species name”, and “Value contains unsupported characters”.

#### Updating taxonomy (‘dataset_update_taxonomy’)

The third step in the build process uses the ‘taxon_list.csv’ configuration file to add additional columns to the taxa table and, if the information is available in the taxon list, to update taxonomic names to the name of the currently accepted/valid taxon concepts. If the ‘aligned_name’ (might include out-of-date taxonomy) and ‘taxon_name’ (currently accepted/valid taxonomy) in the ‘taxon_list.csv’ file differ, the ‘aligned_name’ is then replaced by the ‘taxon_name’ ensuring the most current taxonomy is used throughout the database.

#### Combining sources (‘build_combine’)

The final step in the build process combines the output for the individual datasets into a single database. The database is rebuilt each time a new dataset is added to the database, a change is made to one of the configuration files, or even if a minor change is made to a dataset’s ‘metadata.yml’ or ‘data.csv’ files.

### Tests

Two types of tests have been incorporated into the {traits.build} pipeline using the R package {testthat}: 1) tests to ensure that each dataset’s ‘data.csv’ file, ‘metadata.yml’ file and output comply with expectations; and 2) tests to ensure that the package behaves as described and expected.

The first set of tests checks that metadata files have been correctly filled in and match the corresponding data.csv files. These occur when the database curator runs the function ‘dataset_test()’ on one or more datasets. These tests check that the ‘custom_R_code’ works, essential metadata fields are not missing, metadata values and fields have not been duplicated, and that output tables can be built and converted to and from wide format. The function incorporates custom expectations built from {testthat} and a ‘reporter’ which summarises test passes and fails. These tests quickly identify changes required to the ‘metadata.yml’ file, such as if a column of data has not been properly matched with a trait from the trait dictionary or categorical trait values require substitutions.

The second type of tests uses the unit testing framework of {testthat}, where tests are written and stored in the ‘tests/testthat’ folder and run any time the {traits.build} package is checked (R CMD check) and re-built as part of the R package development workflow. We wrote tests that check that package functions run without error and output is as expected. Within the ‘tests/testthat’ folder we also created several ‘test datasets’ and their expected {traits.build} output tables. During testing, each ‘test dataset’ is rebuilt using the current {traits.build} workflow and the output is compared to the expected output tables; any discrepancies caused by flaws in the code are subsequently caught. These are automated to run on open Pull Requests using GitHub Actions, such that any changes to code that alter the output tables can be identified before merging.

### Reports and error checking

In addition to the automated tests, the {traits.build} workflow includes a function to create an HTML-formatted report for each dataset, ‘dataset_report()’. The report summarises the dataset’s metadata, tabulates categorical trait values, plots numeric trait values against values for the specific trait recorded for other datasets in the database, and summarises taxon names that appear in the dataset. These reports should be reviewed by both the dataset curator and the dataset contributor. Our experience is that it is rare for no errors to emerge during the initial addition of a new dataset and that the reports are an efficient, effective method to quickly detect and rectify errors. The most common error is the mis-mapping of units.

## Discussion

Trait databases have proliferated over the past decade, as large data compilations are increasingly sought for ecological analyses (e.g. https://opentraits.org/datasets.html). As the eventual databases are still small in size by today’s “big data” standards, the primary challenge for trait databases lies with harmonising these diverse small data fragments, rather than engineering challenges of storing large amounts of machine collected data. Indeed, these databases tend to be compiled by individual research groups with expertise in trait ecology, not database standards. To be effective long-term, these researchers must establish both a robust database compilation workflow and, hopefully, develop insightful database documentation, both of which divert time away from their primary goal of amalgamating trait data for analysis. The {traits.build} R package offers a solution by providing an off-the-shelf workflow for building an ecological trait database that can accommodate diverse datasets, can document ecological metadata, has been tested on more than 500 datasets, includes a published data model, and is supported by a series of tutorials and help files.

Acknowledging the dispersed nature of ecological trait data compilations, the Open Traits Network community offered a vision for achieving greater data integration through six Open Science principles (Gallagher et al., 2020). Two of their principles relate directly to database construction, namely ‘Open Resources’ and ‘Open Source’, encouraging researchers to build databases using protocols that are open source, reliable, and error-checked at all stages in database development. To adhere to these principles, the code that harmonises individual datasets into a database must satisfy the following requirements. It must reside in an open repository and be sufficiently documented such that database users can be confident that data from multiple datasets are accurately merged into the output database. It should be generalisable, such that components can be reused to build additional databases, advancing the integration across databases; {traits.build} fulfills these requirements. Moreover, because a database built using the {traits.build} is underpinned by a published data standard, its data content will be more interoperable with other trait databases, increasing its reuse and exposure (Figure 6) (Gallagher et al., 2020; Wilkinson et al., 2016).

**Figure 6.**
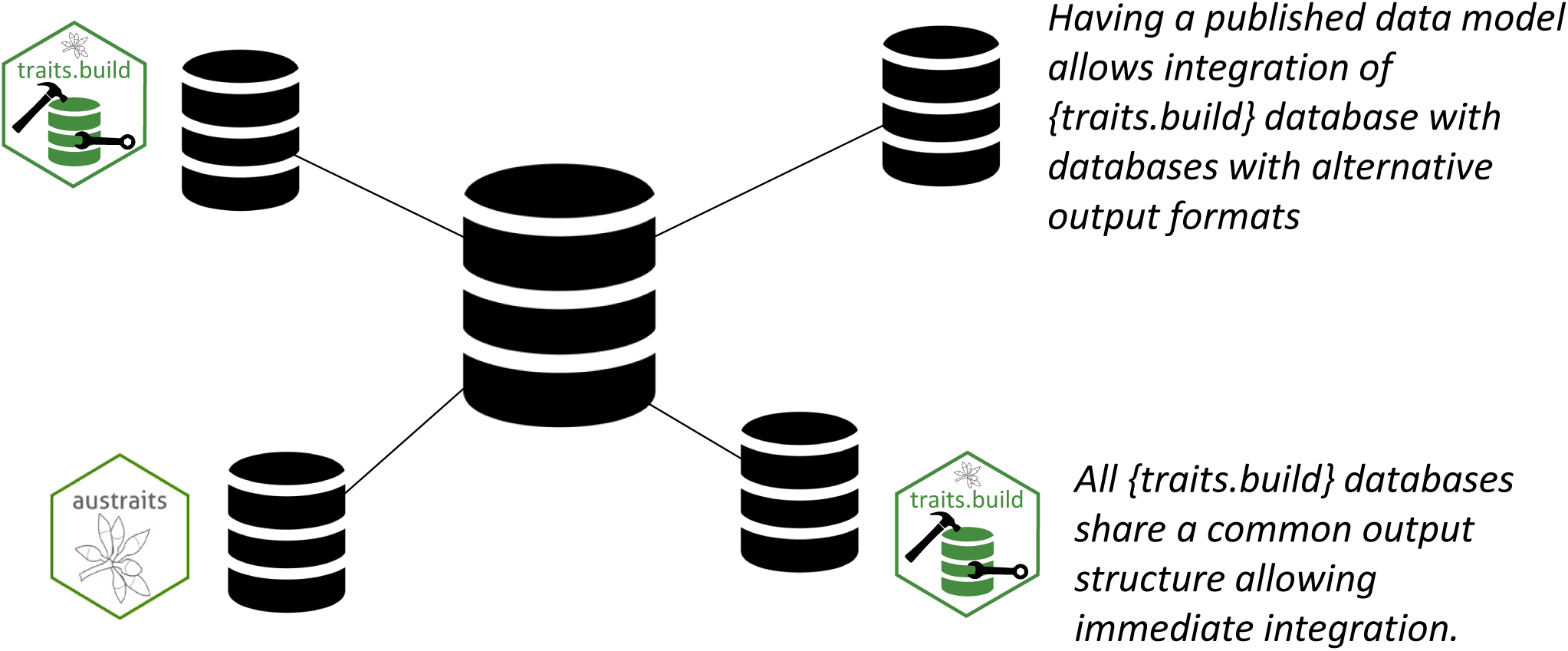
Publishing a data model that includes a description of each column in a {traits.build} databases’ output tables, promotes database integration as envisioned by the Open Traits Network framework (Gallagher 2020). Any database that use the traits.build workflow, creates a database that can automatically be merged with other {trait.build} databases. In addition, the explicit descriptions of each column and mappings to published vocabularies, allow easy integration with other databases.

### Making data FAIRer

The FAIR data principles were proposed to guide knowledge discovery and reuse (Wilkinson et al., 2016). Workflows to make data Findable, Accessible, Interoperable, and Reusable quickly became the expected standard for all databases, but data managers note that these terms are often applied too loosely (Jacobsen et al., 2020; Mons et al., 2020). The {traits.build} workflow strives to adhere to these principles: by publishing the data model as a formal RDF ontology through the w3id.org/traits.build namespace, with permanent resolvable identifiers and rich metadata linked to each term, the database structure is Findable. The metadata are Accessible and the entire {traits.build} workflow is readily accessible on the project’s GitHub repository; each process in the building of a database is transparent. In the strict sense of the FAIR guiding principles, the {traits.build} database structure is Interoperable because each term (both the column headers and other concepts) are mapped to identical or similar terms in other published vocabularies, facilitating the linkage of a {traits.build} database to other trait databases. Furthermore, by offering a workflow that can output multiple databases with identical structures, the {traits.build} package is allowing the creation of entirely interoperable databases. Finally, the data resources will be fully Reusable if the database metadata encapsulated within the databases ‘metadata.yml’ file is included in public database releases; for instance, the completed database should be shared via a repository such as Zenodo, using a CC BY 4.0 license and ensuring project metadata is included, especially that the database is compiled by ‘https://github.com/traitecoevo/traits.build/’.

### Extending existing resources

The need for both established workflows and established database standards for ecological trait databases has been recognised for many years, with an increase in solutions presented over the past decade (Kattge et al., 2011; Lenters et al., 2021; Madin et al., 2007; Schneider et al., 2019). The {traits.build} workflow has taken inspiration from these initiatives and, adhering to the vision of the bioinformatics community, strives to create a workflow that extends (not recreates) existing resources, documents its data model in a format simultaneously machine-readable and accessible to human users, and works towards creating a reusable community standard (Fernández-López et al., 2019; Gkoutos, 2006; Kamdar et al., 2017).

The two most recent database standards we reference, ETS (Schneider et al., 2019) and Lenters (2021), were established simultaneously with (and independently to) the {traits.build} standard, yet the three data models overlap substantially. The {traits.build} team is heartened by this, as it indicates an emerging coalescence across research groups in best practice database standards and protocols. The ‘traits.yml’ configuration file required by the {traits.build} workflow includes a dictionary of explicit trait concepts, descriptions, allowable ranges, allowable categorical values, and output units; ETS has identical requirements (Schneider et al., 2019) and a similar reporting format has been proposed for physiological traits (Ely et al., 2021). Having identical requirements facilitates cross-mapping terms between database structures, as the meaning of the terms is truly identical, allowing the corresponding term to be mapped as a “skos:exactMatch”. Meanwhile, while Lenters et al. (2021) does not include a data model it identifies that published trait definitions, metadata forms, a published data model/ontology, taxonomic standardisation and column header standardisation as essential components to compile a trait database, all features integrated into the {traits.build} inputs workflow, and output structure.

### Better mapping of context properties

One key extension of the {traits.build} workflow over existing trait database workflows is the detailed mapping of context properties. The ability to flexibly capture contexts is essential to fully adhere to the OBOE data model. Moreover, interactions with researchers have shown that they may be hesitant to allow their datasets to be integrated into a database if they perceive that rich and essential contextual information will be lost and trait values misinterpreted when used in new analyses (pers. comm. AusTraits data curators). For instance, physiological ecology research typically requires that the conditions under which a trait is measured (i.e., the context) are captured and should be retained together with the relevant trait measurements (Ely et al., 2021). To this end, all contextual information can be accurately documented by the {traits.build} structure, and by dividing context properties into multiple categories, the linkage between context property values and trait observations is made explicit (Figure 1).

### Importance of database curators

While Lenters et al. (2021) and Schneider et al. (2019) describe processes for individual researchers to fill in the metadata for their studies, the experience of the AusTraits, AusInverTraits, and AusFizz databases has been that it is more reliable and efficient for a small team of database curators to propagate metadata files for datasets for several reasons. Firstly, these curators will be familiar with the {traits.build} functions, ontology, and the accompanying trait dictionary, minimizing errors during metadata file construction. Errors can be time-consuming to trouble-shoot and mis-mapping of data columns to the correct trait concept can be difficult for reviewers to detect. Second, the experience of the AusTraits team has been that many researchers enthusiastically contribute their previously published datasets because this only entails submitting a spreadsheet and manuscript DOI. They would not have the research time to commit to learning how to fill in a metadata file and, if this were a requirement, they would simply not submit their datasets.

### Aligning with OBOE

While several ecological trait databases (Kattge et al., 2020; Leinfelder et al., 2011; Madin et al., 2016) have been built around the OBOE structure, a reusable workflow to build OBOE-adherent trait databases has not been previously established. The {traits.build} data standard closely follows the OBOE structure, offering a workflow for researchers to build trait databases for diverse taxonomic groups, trait categories, or geographic regions that is underpinned by a semantic data model. Only a few relational ontologies exist that are appropriate for observational data (SOSA, EML, OBOE), and of these, OBOE is explicitly designed for integration into data models for ecological databases that includes traits, accompanied by context properties and location properties (Madin et al., 2007).

SOSA, the Sensor, Observation, Sample, and Actuator ontology, also offers a semantic structure for documenting information about observations, but is centrally focused on information about the sensor, which is not provided for many ecological datasets (Janowicz et al., 2019). None-the-less cross-mappings between SOSA and OBOE have been established and it would be easy to link a {traits.build} database to one following the SOSA model. The complexity of the EML and large number of essential variables has hindered its use, as it is impractical to fully propagate the requested metadata for this standard (Mena-Garcés et al., 2011).

### Future work

In the spirit of the Open Traits Network vision for database creation in trait science, the {traits.build} data model, output structure and workflow must not only be useful and reliable in its current format, but also be able to be refined following community requests (Figure 6). The process of building a {traits.build} database must be sufficiently intuitive to entice new adopters to use this workflow instead of devising their own. The functions developed to seamlessly populate the dataset ‘metadata.yml’ files and the tutorials and other help materials hosted on the supplemental GitHub repositories both contribute to these goals. The user community can request edits to the existing tutorials or the creation of additional tutorials to enhance their experience. Recent refinements follow suggestions from research groups currently building databases using the {traits.build} workflow. For instance, the research group building the AusInverTraits database requested that ‘entity_type’ be expanded to include trait values that applied to all members of a genus, family or order, rather than just species, populations and individuals. The capability to map in response curve data has been added per request of the plant physiology and animal thermal tolerance communities.

Future additions will include a ‘catalogue number’, to be developed in consultation with the Global Biodiversity Information Facility (GBIF). This will allow observations within a {traits.build} database to be explicitly linked to herbarium vouchers or occurrence data, for instance on iNaturalist, and ensure ‘traits.build’ databases can be integrated into GBIF’s upcoming traits extension. The open source, modular nature of the traits.build workflow means that refinements and additions are straightforward to implement and test, broadening the appeal of the {traits.build} package to ever more situations. With continued support, the {traits.build} training materials could be translated to additional languages, increasing the global uptake of the tools and in-person and online training courses could rapidly introduce new users to {traits.build}. The advances in database development presented here are intended for use across the trait science community. Openly sharing the development of {traits.build} creates greater research and rewards funders for their commitment to database development.

## Acknowledgements

The AusTraits project received investment (https://doi.org/10.47486/DP720) from the Australian Research Data Commons (ARDC). The ARDC is funded by the National Collaborative Research Infrastructure Strategy (NCRIS). We thank the ARDC for their commitment to training Australia’s researchers in best-practise informatics practices via their co-investment strategy, whereby they offer easily-accessed informatics expertise to their grantees.

## Supplemental Tables

**Table S1.**
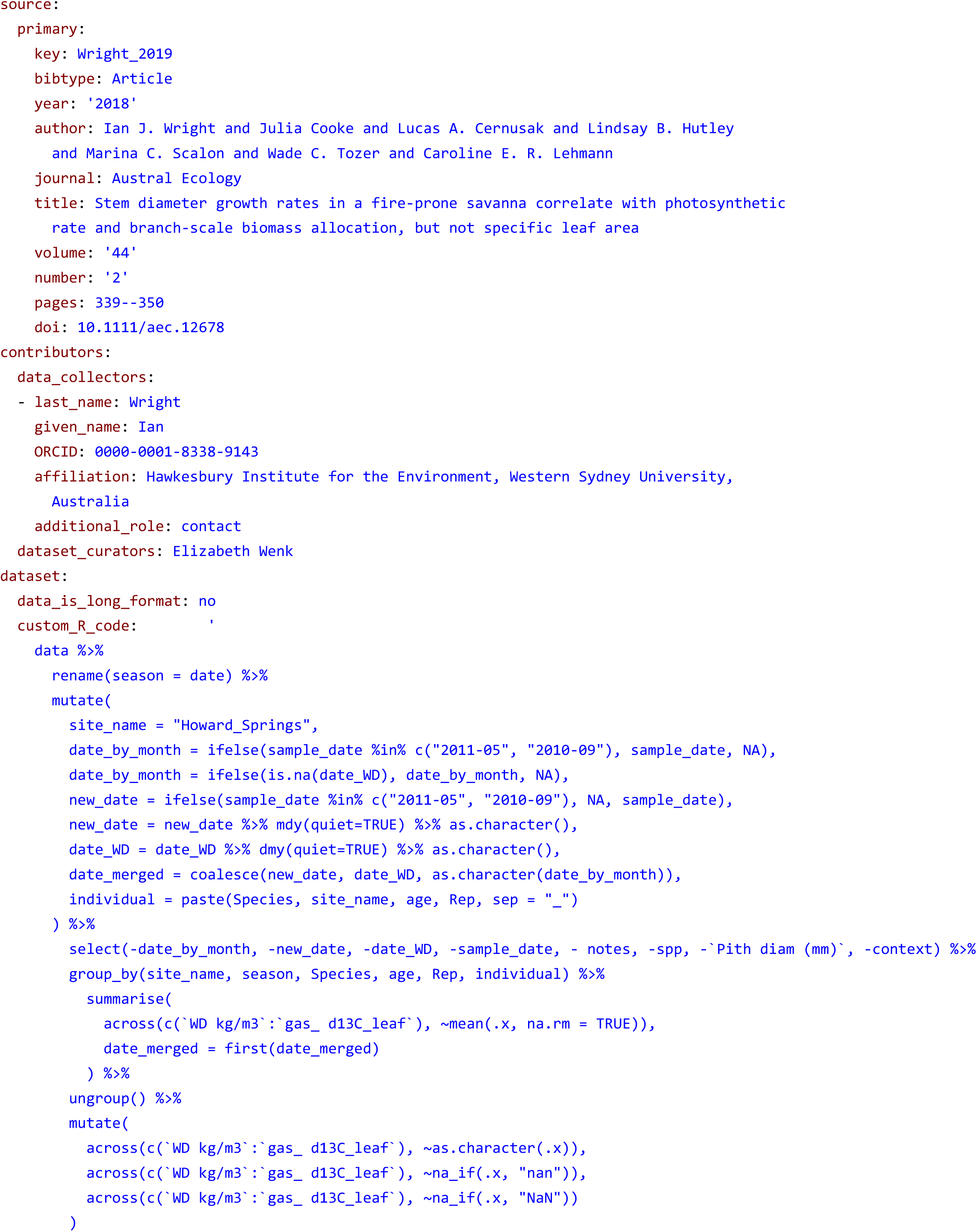
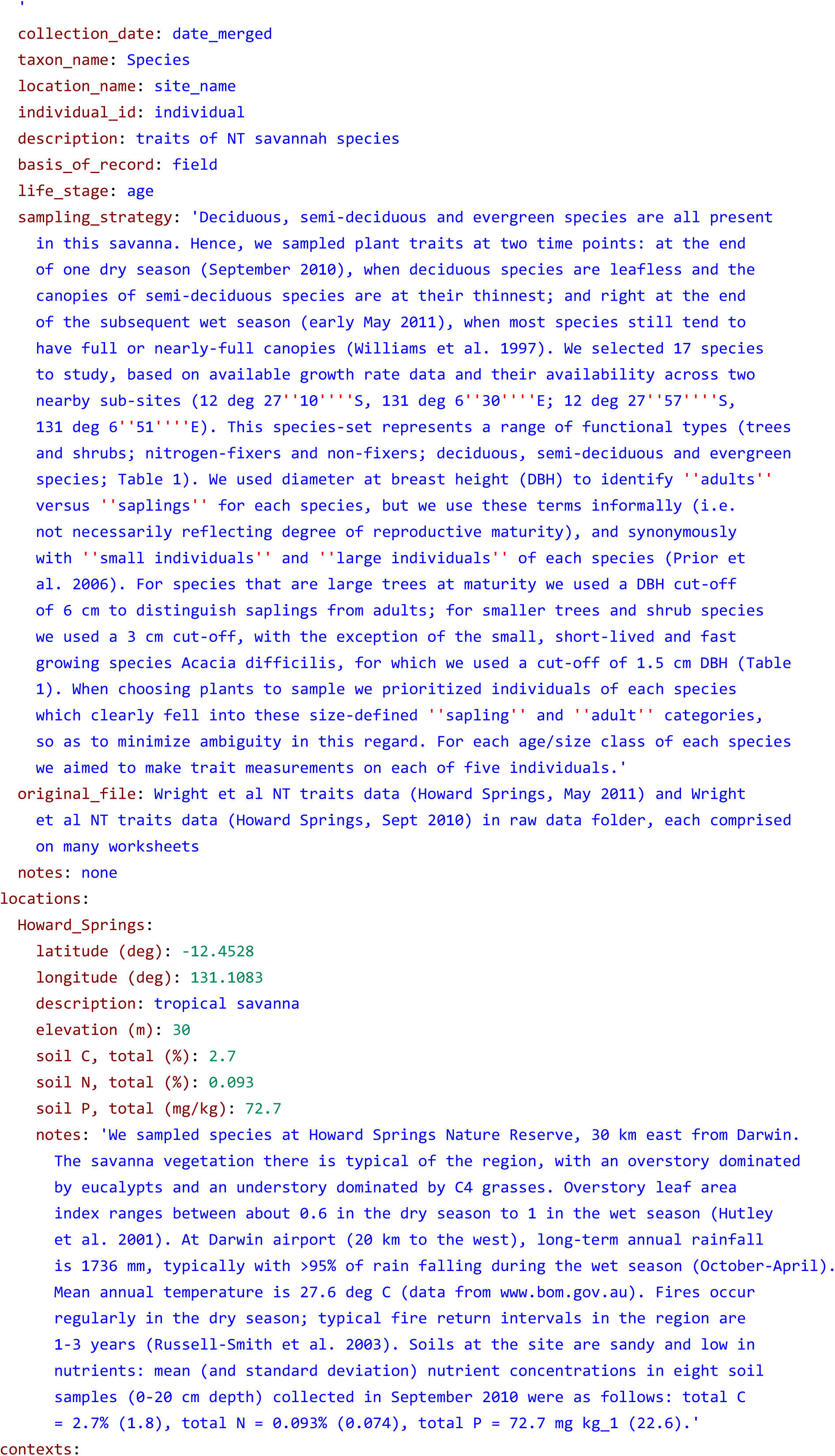
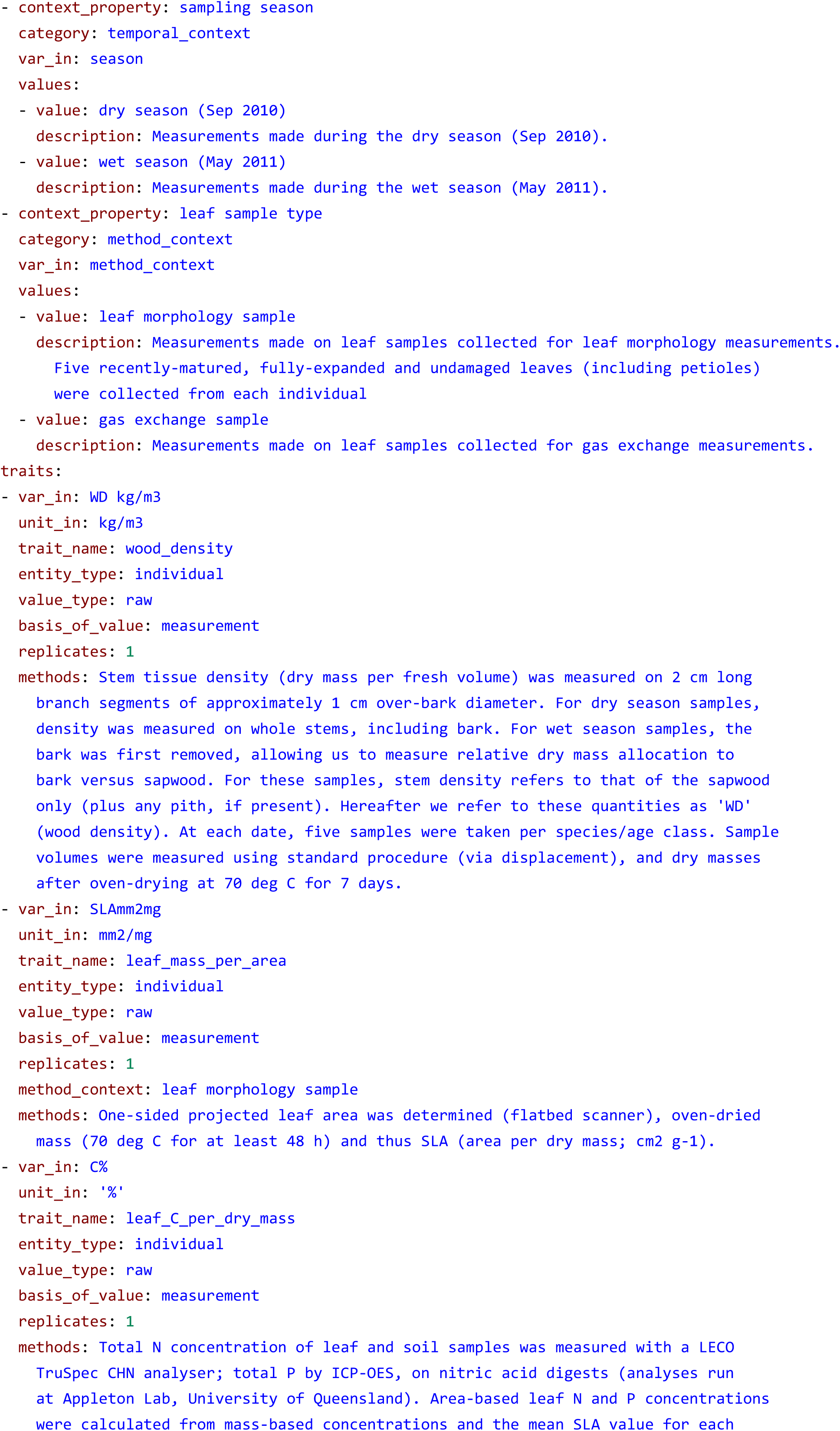
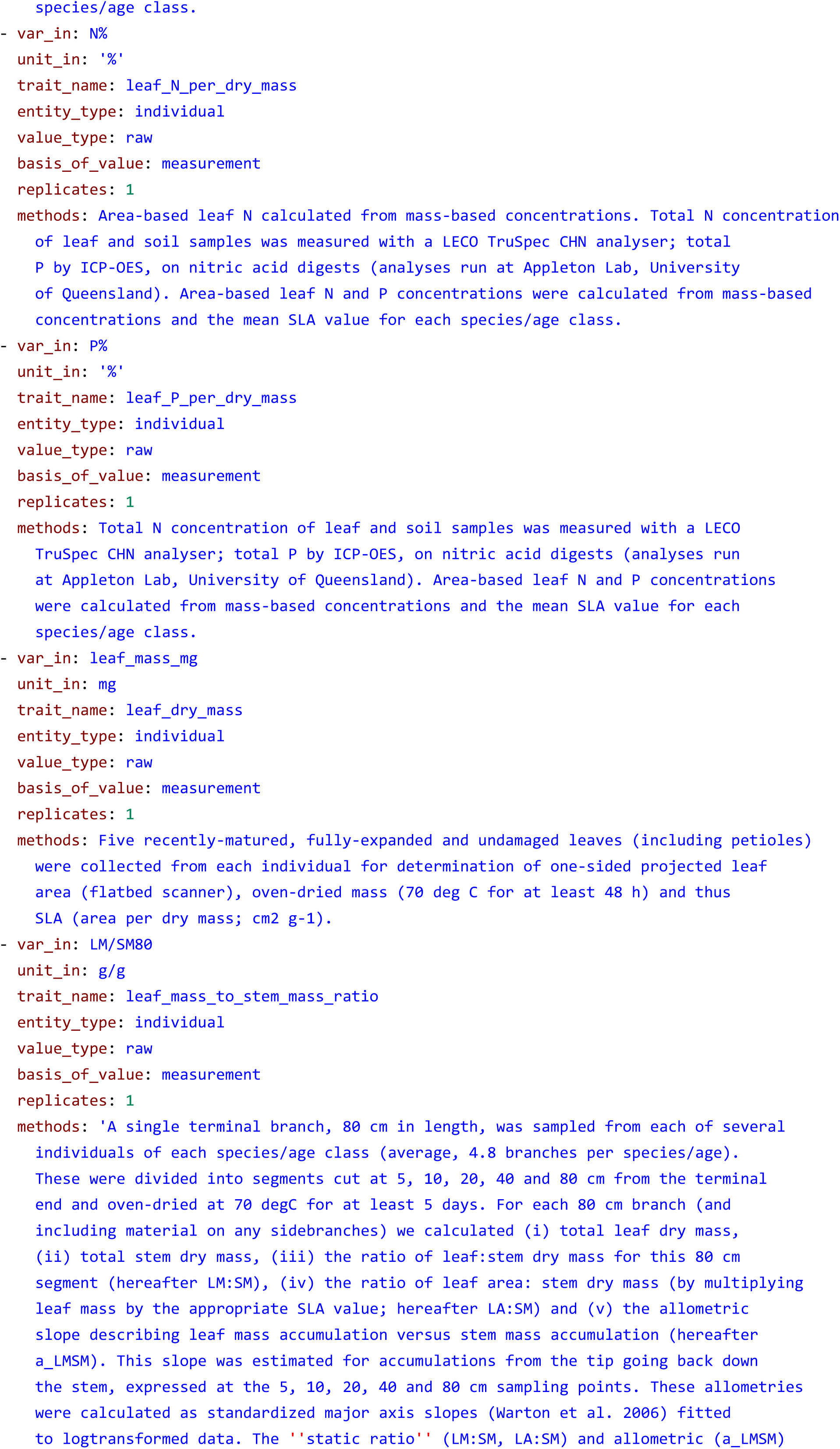
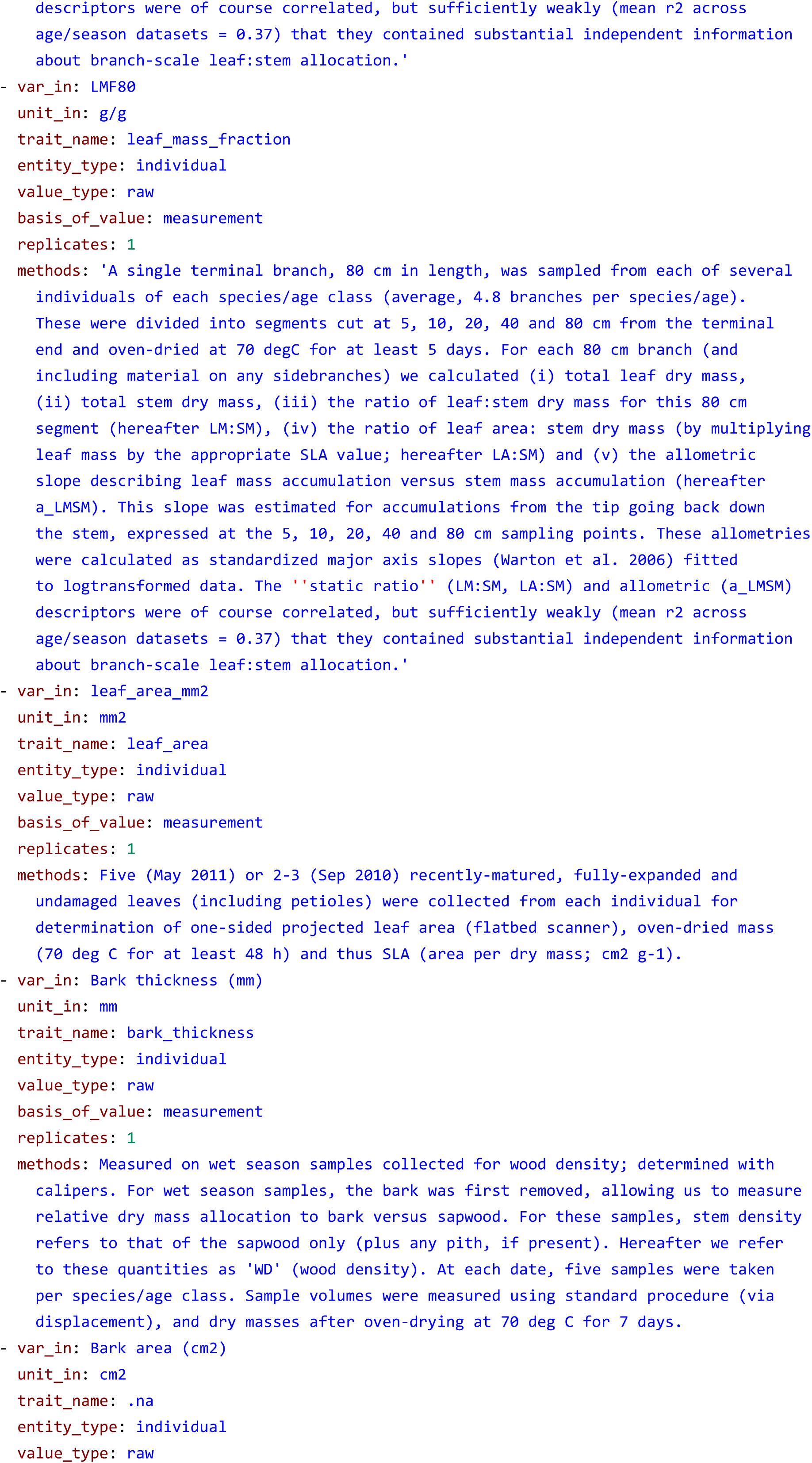
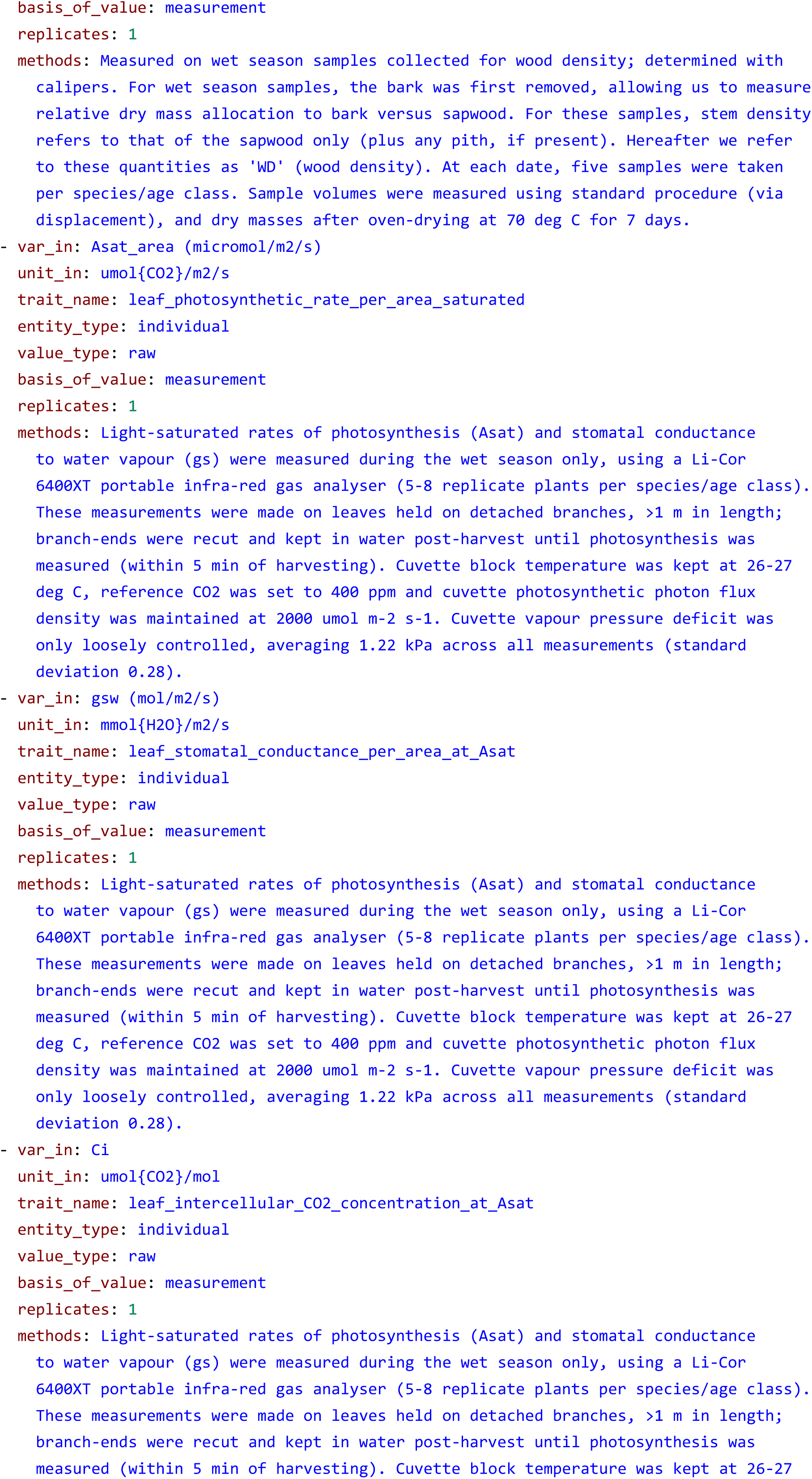
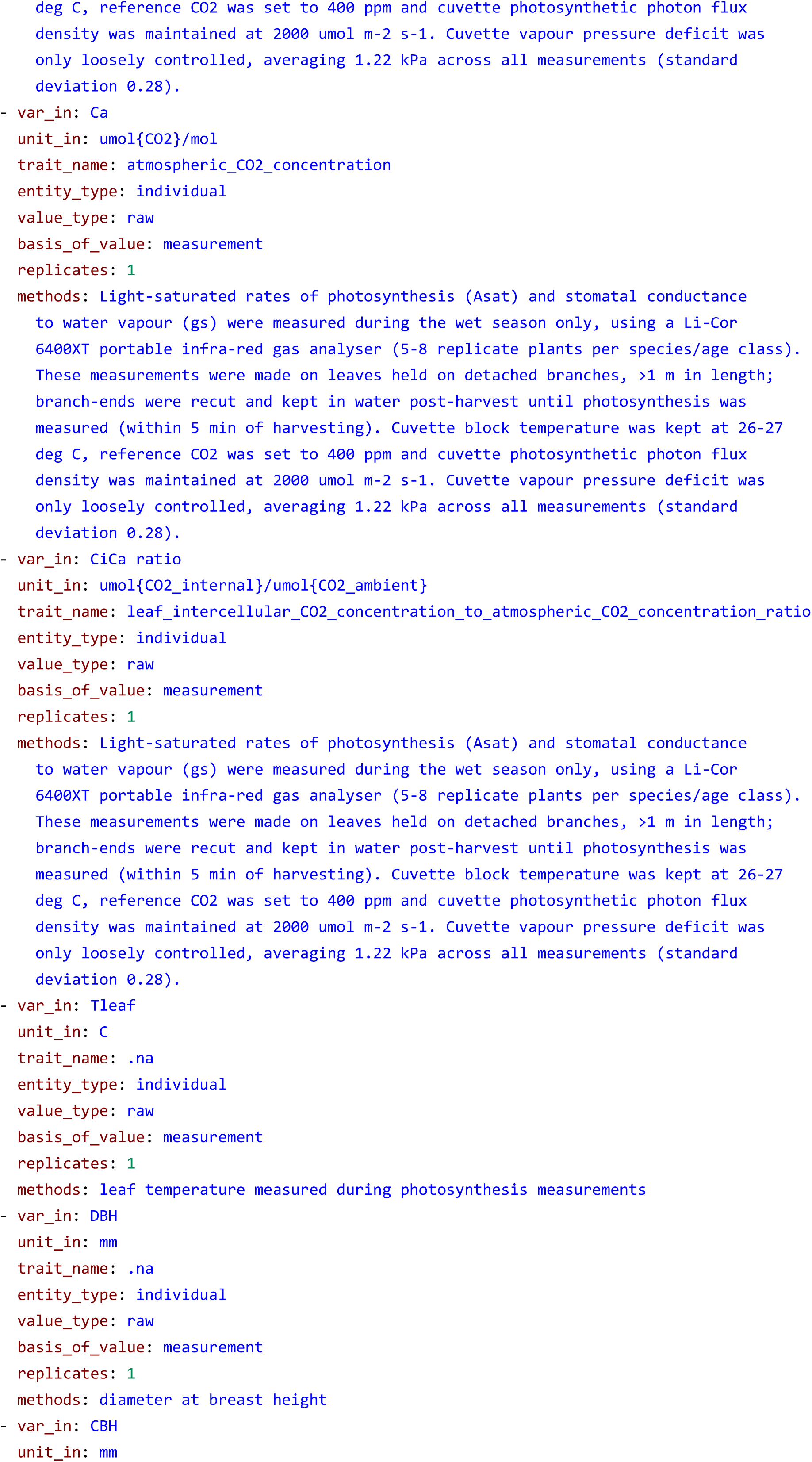
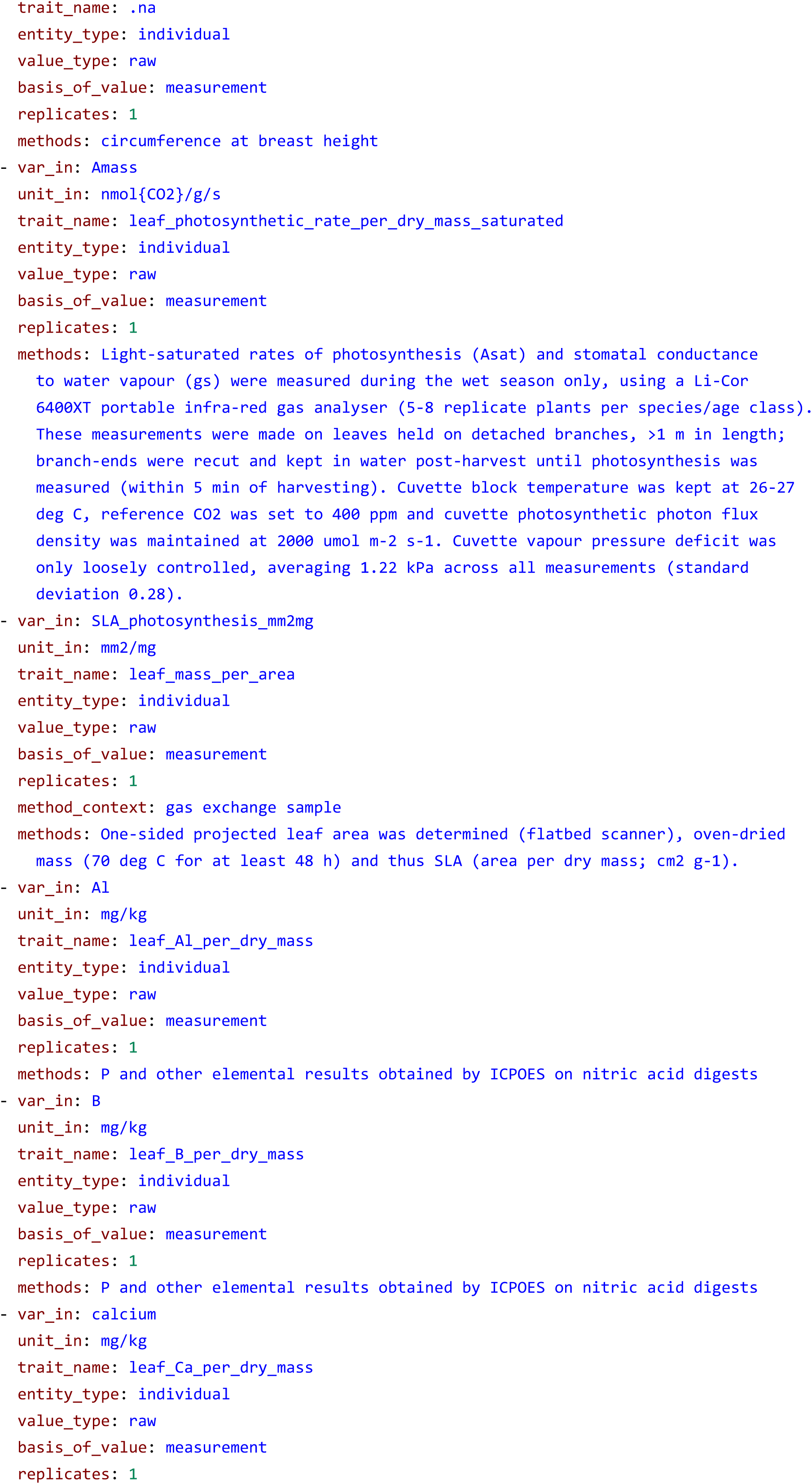
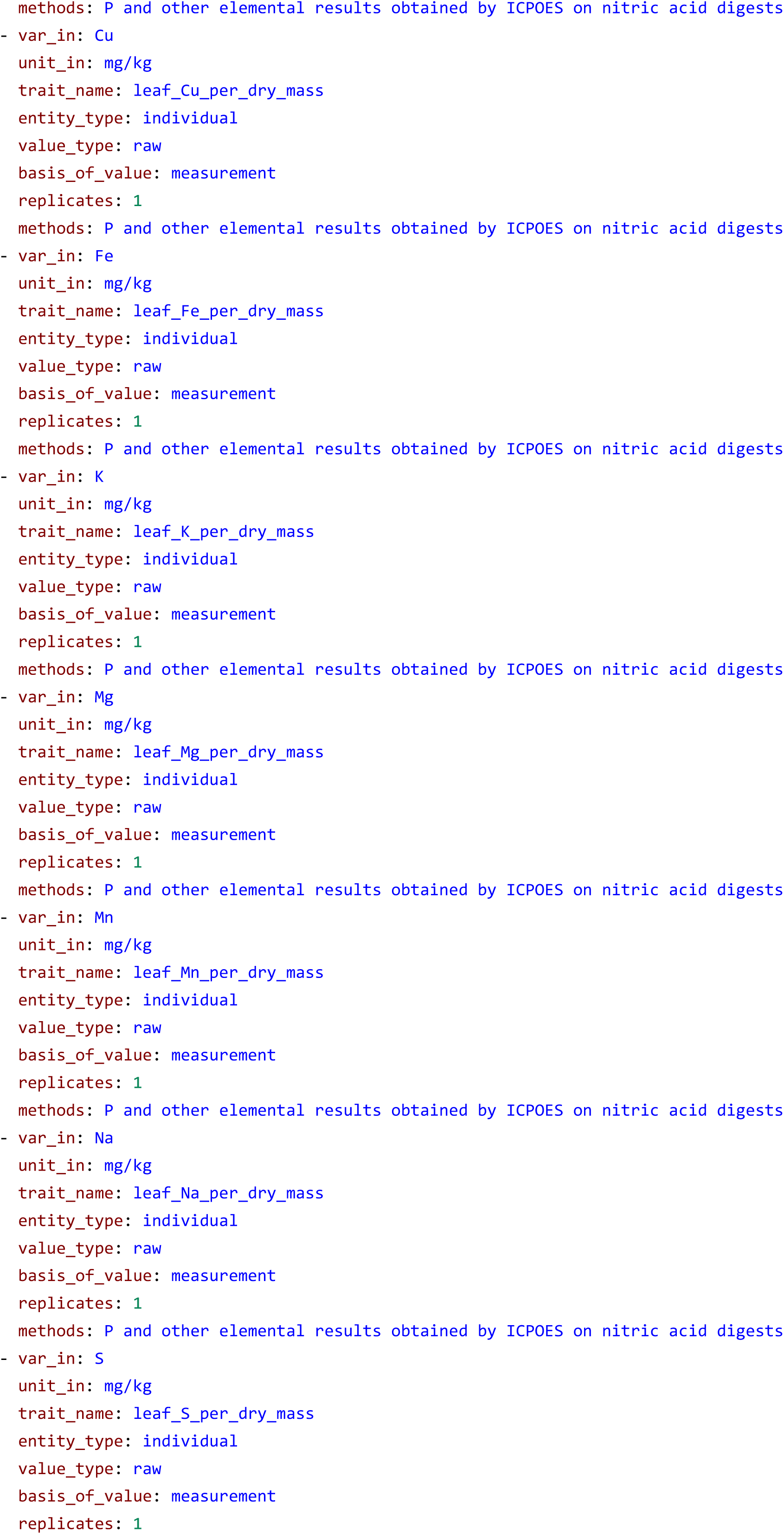
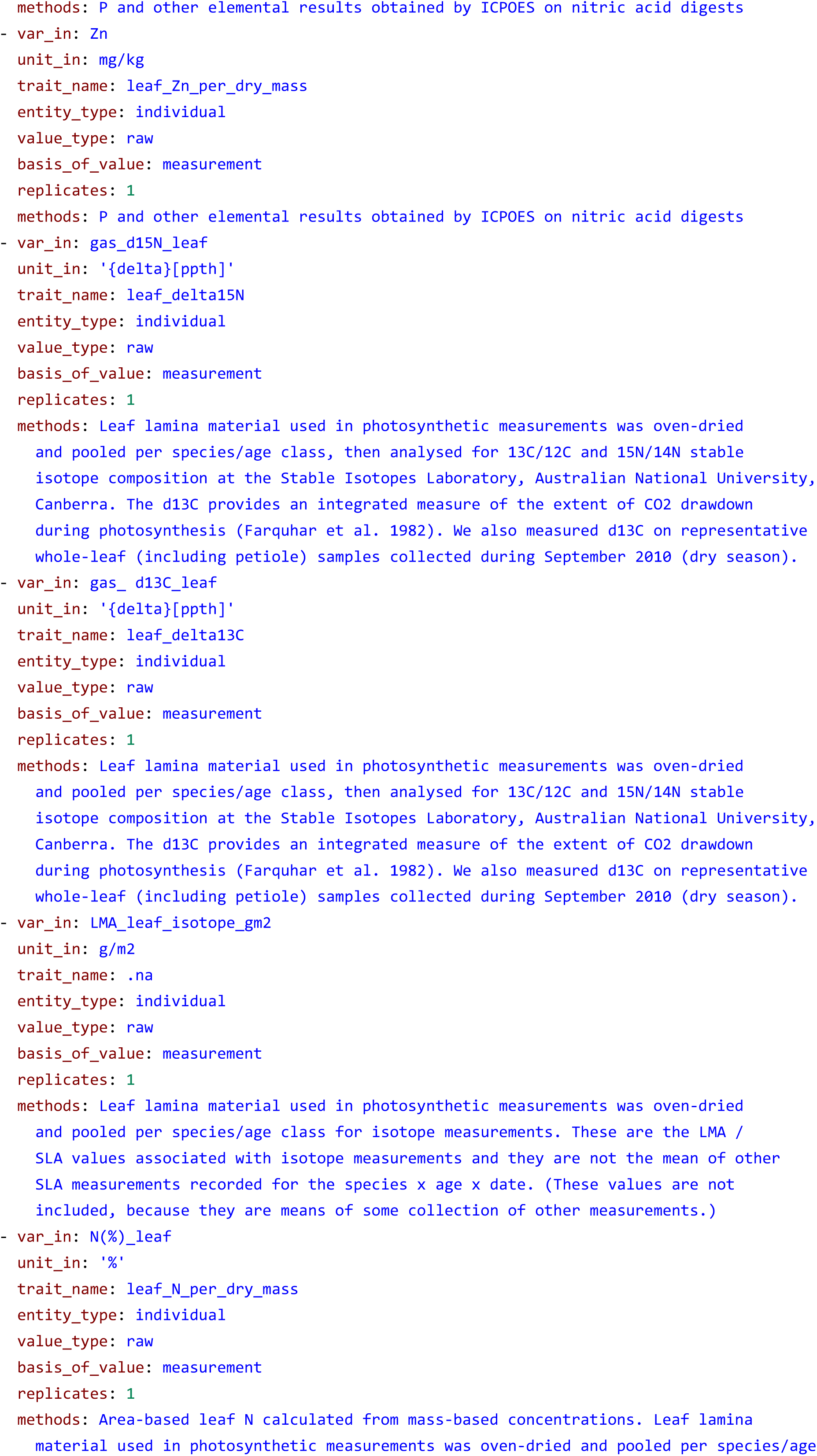
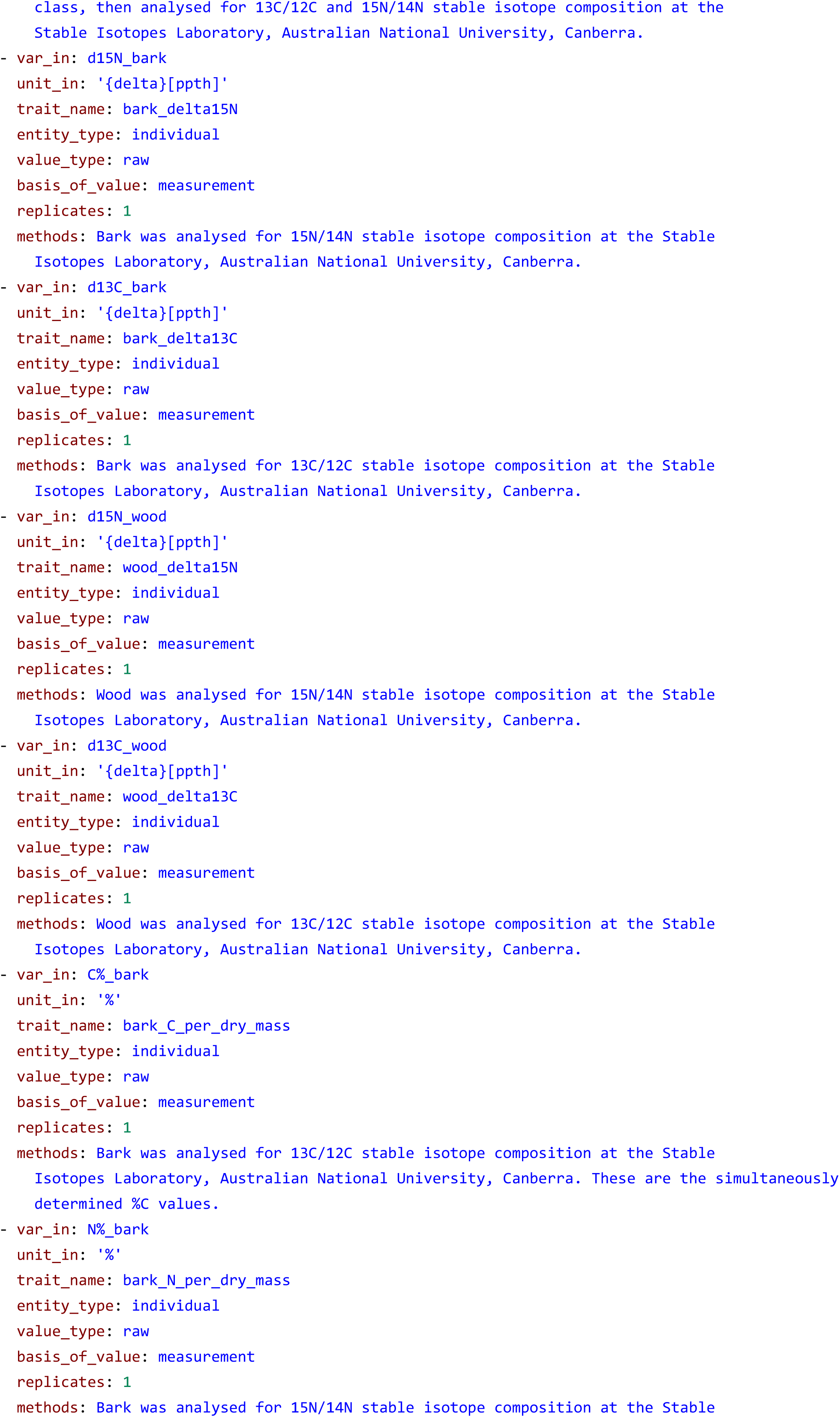
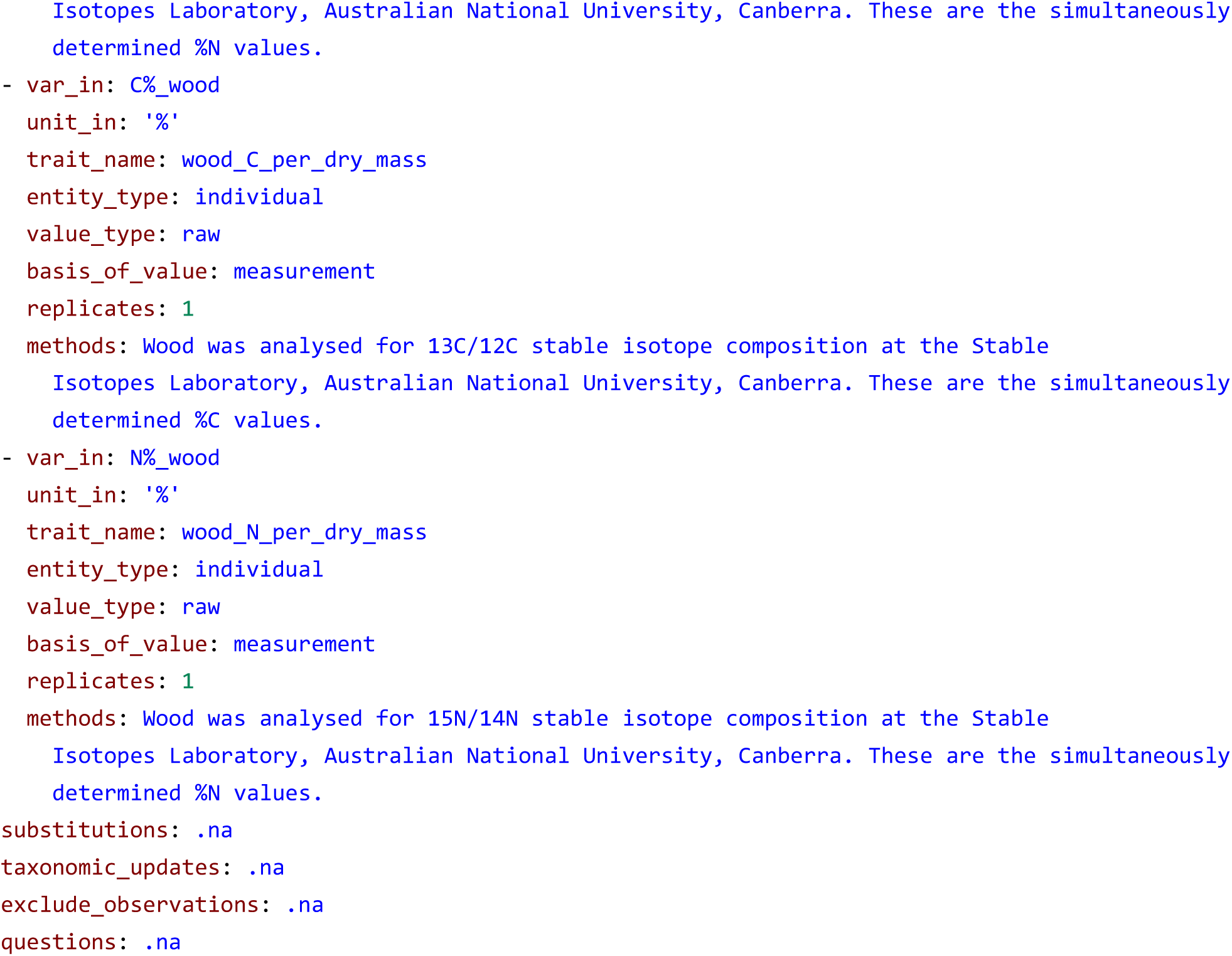
Example of a metadata.yml file from the AusTraits database, available on the project’s Github repository at <u>htps://github.com/traitecoevo/austraits.build/tree/develop/data/Wright_2019/metadata.yml</u>

**Table S2.** Context property categories. Although it is best practice to use the same syntax for the same context property throughout a database, the {traits.build} data model does not require this and similarly does not require a dictionary of defined context properties.

| context property categories | examples | description |
| --- | --- | --- |
| treatment | Nitrogen addition, drought treatment | Measurement made as part of an observation offers context about experimental treatments for a trait-centered observation. |
| plot | Fire history, slope position | Measurement made as part of an observation offers context about plot properties for a trait-centered observation. |
| entity | Sex, growth history | Measurement made on an entity that are themselves not trait measurements, but offer context about the entity for a trait-centered observation. |
| temporal | Sampling season, sampling time of day | Measurement made as part of an observation offers context about a specific point in time a trait-centered observation. |
| method | Canopy position, distance from branch tip sampled | Information that refines a described trait-measurement method, explaining a stratified measurement method. |

**Table S3.** Columns within the locations table and their core mappings in the {traits.build} data model.

| column | description | properties | ontology mapping |
| --- | --- | --- | --- |
| dataset_id | Primary identifier for each study contributed to AusTraits, most often these are scientific papers, books, or online resources. By default this should be the name of the first author and year of publication, e.g. `Falster_2005`. | is unique identifier for `observations_from_dataset` | oboe-core:Identifier |
| location_id | A unique integer identifier for a location, with locations numbered sequentially within a dataset. The identifier links to specific information in the location table. | is unique identifier for `observations_at_location` | oboe-core:Identifier |
| location_name | The location name used by the data contributor or a generic location name designated by AusTraits when one is not provided. | label for `location` | rdfs:label; skos:prefLabel |
| location_property | The location characteristic being recorded. The name should include units of measurement, e.g. `MAT (C)`. Ideally we have at least the following variables for each location, `longitude (deg)`, `latitude (deg)`, `description`. | characteristic for `location`; has value `value` | oboe-core:Characteristic |
| value | The measured value of a location property. |  | oboe-core:MeasuredValue; dwc:measurementValue |

**Table S4.**
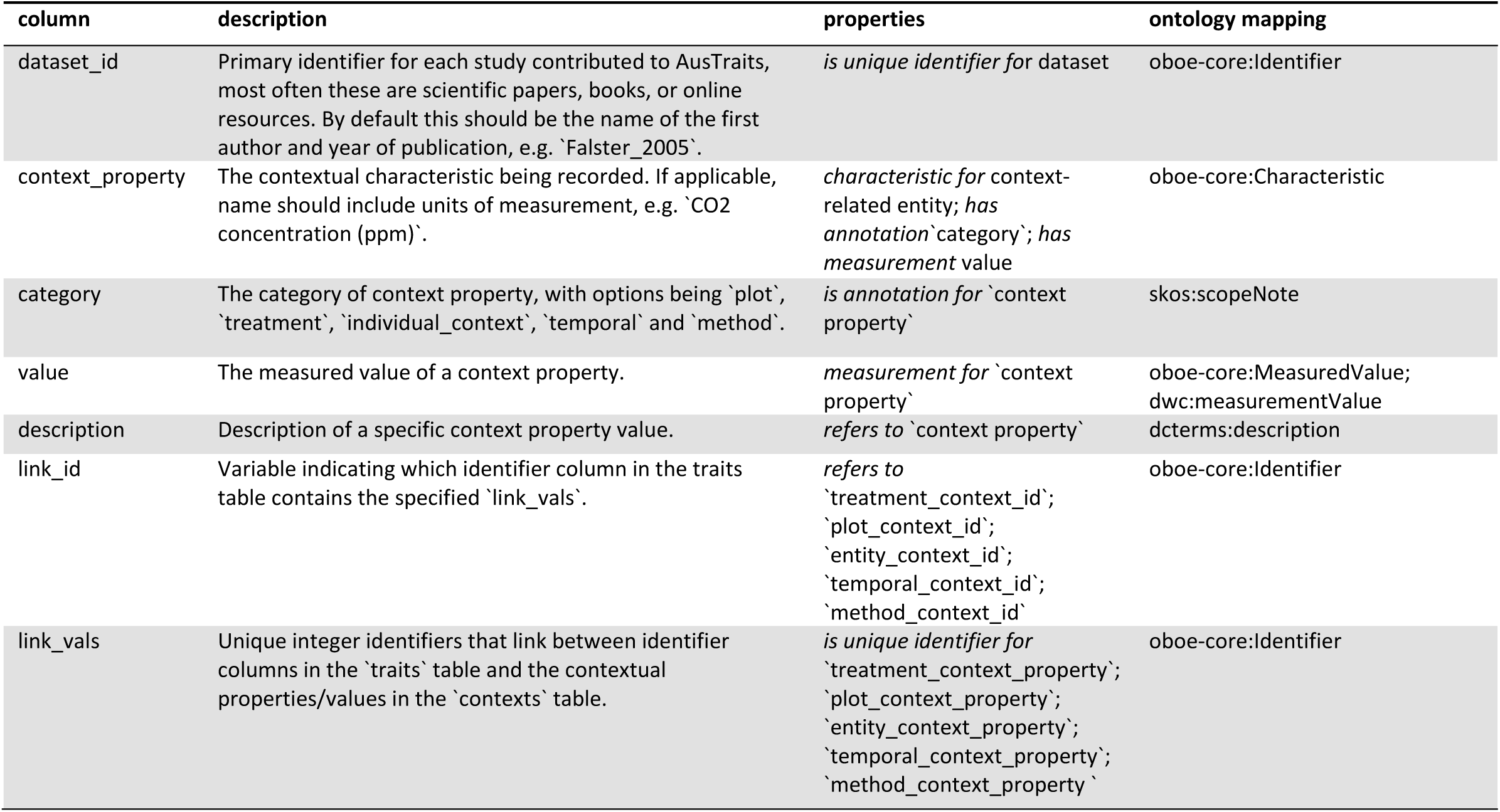
Columns within the contexts table and their core mappings in the {traits.build} data model.

**Table S5.** Columns within the methods table and their core mappings in the {traits.build} data model.

| column | description | properties | ontology mapping |
| --- | --- | --- | --- |
| dataset_id | Primary identifier for each study contributed to AusTraits, most often these are scientific papers, books, or online resources. By default this should be the name of the first author and year of publication, e.g. `Falster_2005`. | *is unique identifier for*<br>`observations_from_dataset` | oboe-core:Identifier |
| trait_name | Name of the trait sampled. Allowable values specified in the table `definitions`. |  |  |
| method_id | A unique integer identifier to distinguish between multiple sets of methods used to measure a single trait within the same dataset. The identifier links to specific information in the methods table. | *is specialisation of* `methods` | oboe-core:Identifier |
| methods | A textual description of the methods used to collect the trait data. Whenever available, methods are taken near-verbatim from the referenced source. Methods can include descriptions such as 'measured on botanical collections', 'data from the literature', or a detailed description of the field or lab methods used to collect the data. |  | oboe-core:Protocol |
| description | A 1-2 sentence description of the purpose of the study. | *refers to* `methods` | dcterms:description |
| sampling_strategy | When available, this information is lifted verbatim from a published manuscript. For preserved specimens, this field ideally indicates which records were 'sampled' to measure a specific trait. | *refers to*<br>`observations_from_dataset` |  |
| source_primary_key | Citation key for the primary source in `sources`. The key is typically formatted as `Surname_year`. | *is unique identifier for*<br>`source_primary_citation` | oboe-core:Identifier |
| source_primary_citation | Citation for the primary source. This detail is generated from the primary source in the metadata. | *has unique identifier*<br>`source_primary_key`; Cites<br>`dataset` | dcterms:bibliographic citation |
| source_secondary_key | Citation key for the secondary source in `sources`. The key is typically formatted as `Surname_year`. | *is unique identifier for*<br>`source_secondary_citation` | oboe-core:Identifier |
| source_secondary_citation | Citations for the secondary source. This detail is generated from the secondary source in the metadata. | *has unique identifier*<br>`source_secondary_key`; Cites<br>`dataset` | dcterms:bibliographic citation |
| source_original_dataset_key | Citation key for the original dataset_id in sources, for compilations. The key is typically formatted as `Surname_year`. | *is unique identifier for*<br>`dataset_key` | oboe-core:Identifier |
| source_original_dataset_citation | Citations for the original dataset_id in sources, for compilation. This detail is generated from the original source in the metadata. | *has unique identifier*<br>`source_original_dataset_key`; Cites<br>`source` | dcterms:bibliographic citation |
| contributor | A person contributing to a dataset. | *refers to*<br>`observations_from_dataset` |  |
| assistants | People who played a more minor role in data collection for the study. | *is specialisation of*<br>`contributor` | dcterms:contributor |
| dataset_curators | Database team member(s) who contacted the data collectors and added the study to the database repository. | *is specialisation of*<br>`contributor` | dcterms:contributor |
| data_collector | Person contributing to a dataset as a data collector. | *is specialisation of*<br>`contributor`; *has unique identifier* ORCID;<br>`given_name`; `last_name`; *has annotation* `affiliation`;<br>`additional_role` | dcterms:contributor |

**Table S6.** Columns within the contributors table and their core mappings in the {traits.build} data model.

| column | description | properties | ontology mapping |
| --- | --- | --- | --- |
| dataset_id | Primary identifier for each study contributed to AusTraits, most often these are scientific papers, books, or online resources. By default this should be the name of the first author and year of publication, e.g. `Falster_2005`. | *is unique identifier for* `observations_from_dataset` | oboe-core:Identifier |
| contributor | A person contributing to a dataset. | *refers to* `observations_from_dataset` | dcterm:contributor |
| data_collector | Person contributing to a dataset as a data collector. | *has unique identifier* `ORCID; given_name; last_name`; *has annotation* `affiliation`; *additional_role`; is specialisation of* `contributor` | gdmt:DataCollector |
| last_name | Last name of the data collector. | *is unique identifier for* `contributor` | obo:IAO_0020017; oboe-core:Identifier |
| given_name | Given names of the data collector. | *is unique identifier for* `contributor` | obo:IAO_0020016; oboe-core:Identifier |
| ORCID | ORCID (Open Researcher and Contributor ID) for the data collector, if available. | *is unique identifier for* `contributor` | obo:IAO_0000708; oboe-core:Identifier |
| affiliation | Affiliation of data collector. | *is annotation of* `contributor` | gdmt:hasContributorAffiliation |
| additional_role | Any additional roles the data collector had in the study, a field most frequently used to identify which data contributor is the contact person for the dataset. | *is annotation of* `contributor` | gdmt:ContributorRole |

**Table S7.** Columns within the taxa table and their core mappings in the {traits.build} data model.

| column | description | properties | ontology mapping |
| --- | --- | --- | --- |
| taxon_name | Scientific name of the taxon on which traits were sampled, without authorship. When possible, this is the currently accepted (botanical) or valid (zoological) scientific name, but might also be a higher taxonomic level. | *has unique identifier* `taxon_id`; *is unique identifier for* `observations_of_taxon`; *has property* `taxonomic_dataset`; `taxon_rank`; `taxonomic_status`; *has basis* `aligned_name`; `original_name` | rdfs:label; skos:prefLabel; dwc:acceptedNameUsage (close match) |
| original_name | Name given to taxon in the original data supplied by the authors. | *is identifier for* `taxon`; *is base of* `aligned_name`; `taxon_name` | rdfs:label; skos:prefLabel |
| taxonomic_dataset | Name of the taxonomy (tree) that contains this concept. ie. APC, AusMoss etc. | *is property of* `taxon_name` | dwc:nomenclaturalCode |
| taxon_rank | The taxonomic rank of the most specific name in the scientific name. | *is property of* `taxon_name` | dwc:taxonRank |
| trinomial | The infraspecific taxon name match for an original name. This column is assigned `na` for taxon name that are at a broader taxonomic_resolution. | *is identifier for* `taxon` | rdfs:label; skos:prefLabel; dwc:infraspecificEpithet (related match) |
| binomial | The species-level taxon name match for an original name. This column is assigned `na` for taxon name that are at a broader taxonomic_resolution. | *is identifier for* `taxon` | rdfs:label; skos:prefLabel; dwc:specificEpithet (related match) |
| genus | Genus of the taxon without authorship. | *is identifier for* `taxon` | dwc:genus; rdfs:label; skos:prefLabel |
| family | Family of the taxon. | *is identifier for* `taxon` | dwc:family; rdfs:label; skos:prefLabel |
| taxon_distribution | Known distribution of the taxon, by Australian state. | *is property of* `taxon` |  |
| establishment_means | Statement about whether an organism or organisms have been introduced to a given place and time through the direct or indirect activity of modern humans. | *is property of* `taxon` | dwc:establishmentMeans |
| taxonomic_status | The status of the use of the scientificName as a label for the taxon in regard to the 'accepted (or valid) taxonomy'. The assigned taxonomic status must be linked to a specific taxonomic reference that defines the concept. | *is property of* `taxon_name` | dwc:taxonomicStatus |
| scientific_name | The full scientific name, with authorship and date information if known. | *is unique identifier for* `Taxon`; *has unique identifier* `scientific_name_id` | dwc:scientificName |
| taxon_id | An identifier for the set of taxon information (data associated with the taxon class). May be a global unique identifier or an identifier specific to the data set. Must be resolvable within this dataset. | *is unique identifier for* `taxon_name` | dwc:taxonID; oboe-core:Identifier |
| scientific_name_id | An identifier for the set of taxon information (data associated with the taxon class). May be a global unique identifier or an identifier specific to the data set. Must be resolvable within this dataset. | *is unique identifier for* `scientific_name` | dwc:scientificNameID; oboe-core:Identifier |

**Table S8.** Annotation properties used within the ‘traits.build’ vocabulary.

| identifier | label | Entity |
| --- | --- | --- |
| skos:prefLabel | preferred label | http://www.w3.org/2004/02/skos/core#prefLabel |
| rdfs:label | label | http://www.w3.org/2000/01/rdf-schema#label |
| skos:exactMatch | exact match | http://www.w3.org/2004/02/skos/core#exactMatch |
| skos:closeMatch | close match | http://www.w3.org/2004/02/skos/core#closeMatch |
| skos:relatedMatch | related match | http://www.w3.org/2004/02/skos/core#relatedMatch |
| rdfs:subClassOf | subclass of | http://www.w3.org/2000/01/rdf-schema#subClassOf |
| SO:SIO_000602 | computational entity | http://semanticscience.org/resource/SIO_000602 |
| dwcattributes:organizedInClass | organized in class | http://rs.tdwg.org/dwc/terms/attributes/organizedInClass |
| oboe-core:hasContext | has context | http://ecoinformatics.org/oboe/oboe.1.2/oboe-core.owl#hasContext |
| oboe-core:characteristicFor | characteristic for | http://ecoinformatics.org/oboe/oboe.1.2/oboe-core.owl#characteristicFor |
| oboe-core:ofEntity | entity | http://ecoinformatics.org/oboe/oboe.1.2/oboe-core.owl#ofEntity |
| oboe-core:hasValue | has value | http://ecoinformatics.org/oboe/oboe.1.2/oboe-core.owl#hasValue |
| oboe-core:measurementFor | measurement for | http://ecoinformatics.org/oboe/oboe.1.2/oboe-core.owl#measurementFor |
| oboe-core:hasMeasurement | has measurement | http://ecoinformatics.org/oboe/oboe.1.2/oboe-core.owl#hasMeasurement |
| gdmt:Cites | cite | http://vocab.fairdatacollective.org/gdmt/Cites |
| SIO:SIO_000628 | refers to | http://semanticscience.org/resource/SIO_000628 |
| SIO:SIO_000673 | has unique identifier | http://semanticscience.org/resource/SIO_000673 |
| SIO:SIO_000674 | is unique identifier for | http://semanticscience.org/resource/SIO_000674 |
| SIO:SIO_000671 | has identifier | http://semanticscience.org/resource/SIO_000671 |
| SIO:SIO_000672 | is identifier for | http://semanticscience.org/resource/SIO_000672 |
| SIO:SIO_000255 | has annotation | http://semanticscience.org/resource/SIO_000255 |
| SIO:SIO_000254 | is annotation of | http://semanticscience.org/resource/SIO_000254 |
| SIO:SIO_000223 | has property | http://semanticscience.org/resource/SIO_000223 |
| SIO:SIO_000224 | is property of | http://semanticscience.org/resource/SIO_000224 |
| SIO:SIO_001096 | is specialisation of | http://semanticscience.org/resource/SIO_001096 |
| SIO:SIO_000642 | is base for | http://semanticscience.org/resource/SIO_000642 |
| SIO:SIO_000641 | has basis | http://semanticscience.org/resource/SIO_000641 |
| dcterms:description | description | http://purl.org/dc/terms/description |
| rdfs:comment | comment | http://www.w3.org/2000/01/rdf-schema#comment |
| rdfs:datatype | datatype | http://www.w3.org/2000/01/rdf-schema#Datatype |

